# Effect of Dietary Fat on the Metabolism of Energy and Nitrogen, Serum Parameters, Rumen Fermentation, and Microbiota in twin *Hu* Male Lambs

**DOI:** 10.1101/438929

**Authors:** Wenjuan Li, Hui Tao, Naifeng Zhang, Tao Ma, Kaidong Deng, Biao Xie, Peng Jia, Qiyu Diao

## Abstract

**Background:** Fat is the main substance that provides energy to animals. However, the use of fat in twin *Hu* lambs has not been investigated. Thirty pairs of male twin lambs were examined to investigate the effects of dietary fat on the metabolism of energy and nitrogen, ruminal fermentation, and microbial communities. The twins are randomly allotted to two groups (high fat: HF, normal fat: NF). Two diets of equal protein and different fat levels. The metabolism test was made at 50-60 days of age. Nine pairs of twin lambs are slaughtered randomly, and the rumen fluid is collected at 60 days of age.

**Results:** The initial body weight (BW) in the HF group did not differ from that of NF group (*P* > 0.05), but the final BW was tended to higher than that of NF group (0.05 < *P* < 0.1). The digestive energy (DE), metabolism energy (ME), DE/ME in the HF group tend to be higher than those in the NF group (0.05 < *P* < 0.1). Ruminal ammonia nitrogen (NH_3_-N) and the proportion of total volatile fatty acids (TVFA) are higher than that in the NF group (*P* < 0.05). A high throughput sequencing analysis reveals that there were no differences between the two groups in terms of the richness estimates and diversity indices (*P* > 0.05). The *Proteobacteria* and *Fibrobacteres* phyla were higher than that in NF group (*P*<0.05).

**Conclusions:** This study demonstrated that high fat diet before weaning can affect the abundance of several groups of rumen bacteria in rumen, such as significantly increasing phyla *Proteobacteria* and *Fibrobacteres*, and genera of *Succinivibrio*, *Alloprevotella*, and *Saccharofermentans*, but significantly decreasing genera of *Clostridium IV*, *Dialister*, *Roseburia*, and *Butyrivibrio*. And high fat diet improved the performance of lambs at weight gain, energy utilization, and had effect on VFA composition but no effects on serum enzymes and serum hormone.

## Introduction

In human, infant formula is designed to meet all the nutritional needs to promote infant growth and development. Early nutrition has critical effects on the long-term health of adult animals [1,2]. Therefore, it is important to understand the early nutritional regulation of young animals. It is reported that calves fed on an elevated plane of nutrition pre-weaning have higher starter intakes and average daily gain during the weaning period [3].

Fat is an essential nutrient for young animals. It is the main supplier and storage of energy in animals. Fat has been used to feed cattle and sheep for a long time [4–6]. Raeth-Knight [7] suggested that high fat diets can lead to rapid growth, which allows heifers to reach breeding size earlier with lower production costs. Other researchers suggested that fat may be the result of the dietary fat, fat type, additive amounts and interactions.

The influence of dietary fat vary among studies, which could be associated with species of animals, type and concentrations of fat and dietary composition [8,9]. However, little information can be found on the effects of dietary fat on twin *Hu* lambs, a local breed in China and the ewes lamb twice a year with 2-3 lambs in most cases per year. With the similar genetic background in twin lambs, twin *Hu* lambs is an ideal model to use in the present study.

Dietary fat shapes rumen microbiota, most of the study focused on the adult ruminant [10], little is known about the effect of pre-weaning diet on the establishment of the rumen microbiota. Although most of the milk replacer directly went into abomasums due to the closure of the esophageal groove by reflex action, part of the milk leaked into the rumen [11], which becomes substrate with starter for ruminal microbes. We compared high fat diet with normal fat diet to determine the effects of dietary fat on digestion, as well as the metabolism of energy and nitrogen, serum parameters, rumen fermentation and microbiota in pre-ruminant lambs. Then we selected twin lambs as experimental animals in order to highlight the mechanism of diet fat based on a consistent genetic background.

## Materials and Methods

### Ethic approval and consent to participate

The feeding trial was carried out at the Hai-lun sheep Farm (Taizhou City, Jiangsu Province, China). All experiments were conducted according to the Regulations for the Administration of Affairs Concerning Experimental Animals published by the Ministry of Science and Technology, China. The Chinese Academy of Agricultural Sciences Animal Ethics Committee approved all experiments, and humane animal care and handling procedures were followed throughout the experiment.

### Experimental Design, Animal Management and Diet

In this study, 30 pairs of twin *Hu* male lambs at seven days old (BW = 4.22±0.56 kg) were obtained randomly and assigned to two groups within block to 20 pens (pen=3 animals, 10 pens/group). All the lambs were weighed and ear-tagged before the start of the experiment, then they were subjected to normal immunisation procedures. Two diets of equal protein but different fat levels were fed to the lambs until 60 days of age: one was a normal-fat diet consisting of a milk replacer (MR; 15% fat, which followed industry standard ‘milk replacer for lamb NY/T2999-2016’ in China and patent ‘a milk replacer for calf and lamb ZL02128844.5’) and a starter (2.8% fat), and the other was a high-fat diet consisting of MR (27% fat) and a starter (5.07% fat). Table 1 presents the chemical constituents and ingredients of the trial diets. All the lambs were fed MR at 2% of BW from 7 to 50 days of age and 1.5% of BW from 50 to 60 days of age. One-third of the MR was fed at 06:00, one-third was fed at 12:00 and the remainder were fed at 17:30. From day 50 to 60, the lambs were fed twice daily at 6:00 and 17:30, allowing 5-10% orts, and fresh drinking water and starter were provided ad libitum.

**Table 1.**
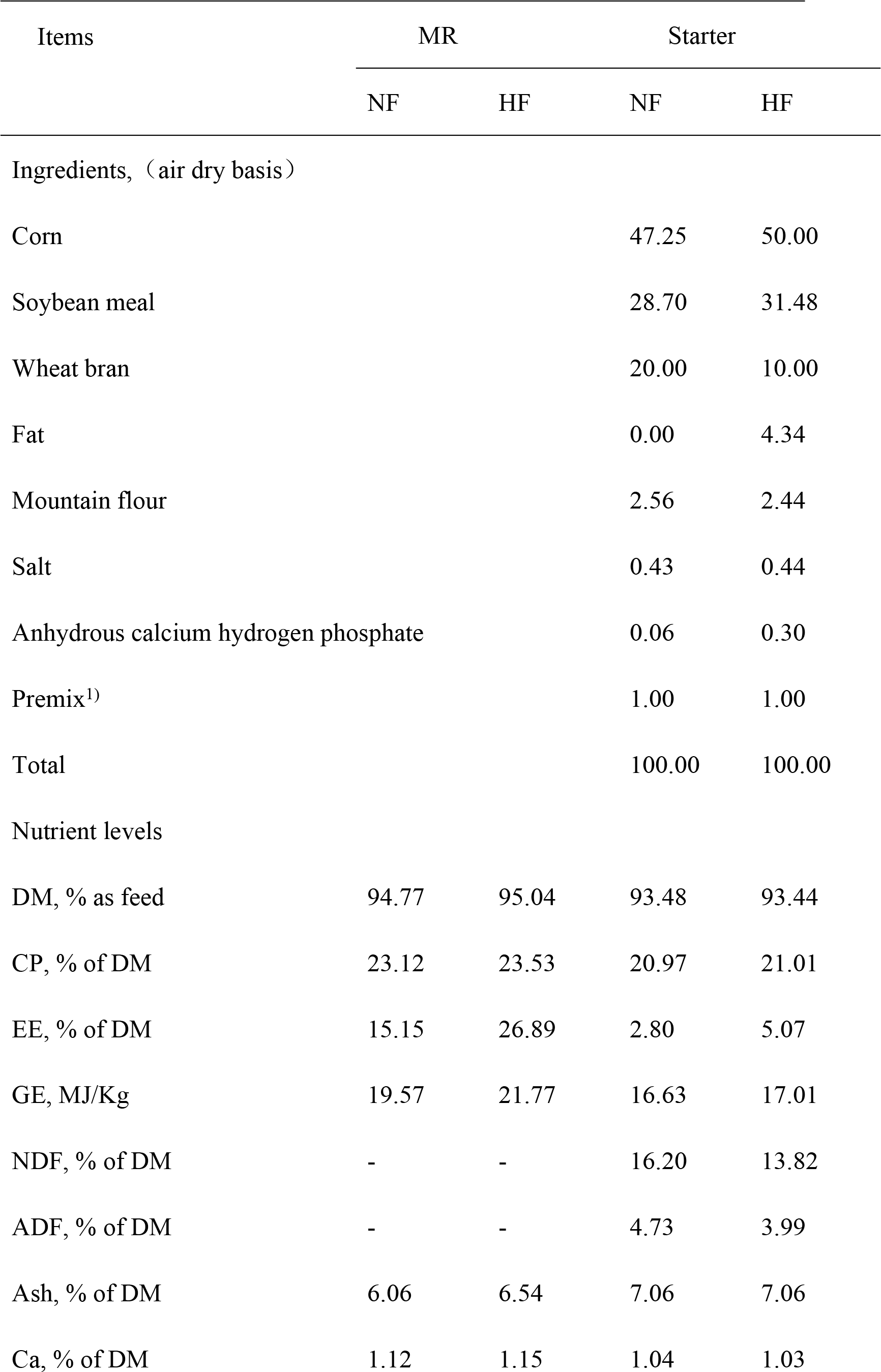
Ingredients and nutritional composition of milk replacer and starter (%).

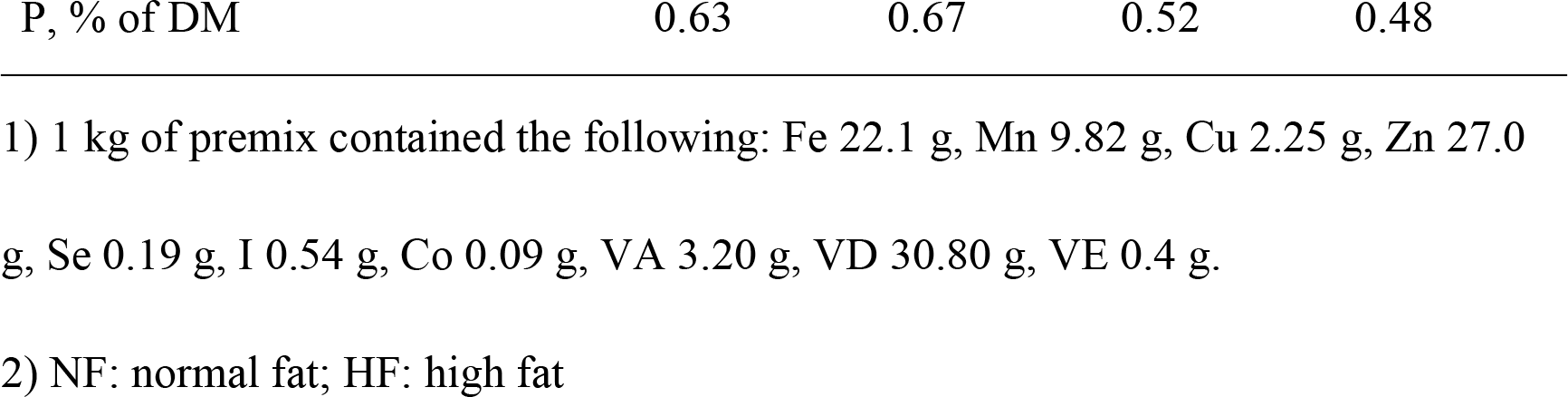

### Sample Collection

From 50-60 days of age, nine pairs of twin lambs were randomly selected according to average weight for digestion and metabolism trail. The feeding period was five days, and the collection period was five days. Faecal and urine samples were collected before morning feeding using total collection sampling and stored at −20 °C for further analysis.

On the 60th day, the lambs which were selected for digestion and metabolism trail were weighed and collected blood via the jugular vein before the morning feeding. The samples were centrifuged at 3,000 × g for 15 min at 4 °C to collect the serum, separated into three aliquots and then frozen at −20 °C for subsequent biochemical index analyses. Nine pairs lambs that were randomly selected by average weight were slaughtered. Feed was withheld for 24 h before slaughtering.

Approximately 150 mL of ruminal fluid sample consisting of a mixture of liquids and solids was obtained. The ruminal fluid pH was measured immediately after collection using a digital pH meter (Testo205 type, Germany). Next, 0.25 mL of metaphosphoric acid (25 g/100 mL) was added to four aliquots of 1 ml rumen fluid, which were centrifuged at 20000 × g at 4 °C for 15 min to determine the VFA and NH3–N concentrations. Two aliquots of 2 mL samples were taken to determine the concentration of microbial protein. Three aliquots of 1.5 mL samples were taken and kept in liquid N for rumen bacterial 16S rRNA analysis.

### Chemical Analysis

Feed, orts and faeces were oven-dried at 65°C for 72 h, smashed using a mill (Wiley, A. H. Thomas Co., Philadelphia, PA, USA) with a 1 mm screen in order to analyse the DM (method 930.15) [12]. The GE of the feed was measured using a bomb calorimeter (6400, PARR Works Inc., USA). The rumen liquid NH_3_–N was measured according to Bremner and Keeney’s [13] method, which calls for the use of a spectrophotometer (UV-6100, Mapada Instruments Co., Ltd., Shanghai, China). The VFA was quantified using a high-performance gas chromatograph (HPGC; GC-128; INESA Corporation) that was equipped with a hydrogen flame detector and a capillary column (FFAP, Zhonghuida Instruments Co., Ltd., Dalian, China; 50 m long, 0.32 mm diameter, 0.50 µm film). The microbial protein was then quantified using the purine derivative method [14].

### DNA Extraction and 16S rRNA Pyrosequencing

First, the sample was melted on ice, and then it was centrifuged and thoroughly mixed. Total cellular DNA was extracted separately using a commercially available kit according to the manufacturer’s instructions (Omega Bio-tek, Norcross, US). DNA concentration and purity were evaluated using a spectrophotometer (Thermo NanoDrop 2000, US). DNA was separated by 1% agarose gel electrophoresis. In the PCR amplification system, each 40 μL PCR mixture consisted of a 10 ng DNA sample, a 1μL forward primer (5 umol), a 4 μL DNA template (0.25 mmol), a 1 μL reverse primer (5 umol) and the 20 μL 2×KAPA HiFi Hotstart ReadyMix. Finally, 40 μL of supplementary sterile water was distilled. Bacterial 16S rRNA genes of the V3-V4 region were then amplified from the extracted DNA using the barcoded primers 341F (5′-ACTCCTACGGGRSGCAGCAG-3′) and 806R (5′ GACTACVVGGGTATCTAATC-3′). The PCR procedure was as follows: 95 °C for 3 min followed by 24 cycles of 98 °C for 15 s, 72 °C for 10 s, 72 °C for 10 s, 94 °C for 20 s, 65 °C for 10 s and 72 °C for 10 s, along with 16 cycles of 94 °C for 20 s, 58 °C for 30 s and 72 °C for 10 s, as well as an extension at 72 °C for 150 s and storage at 4 °C. Amplicons were separated from 2% agarose gels and purified using the AxyPrep DNA Gel Extraction Kit (Axygen Biosciences, Union City, CA, USA). According to the manufacturer’s instructions, the library size was about paired-end reads of 300 bp using the 1% agarose gel test. The purified PCR product library was detected and quantified using the Qubit^®^ dsDNA HS Assay Kit, followed by a corresponding proportion of each sample according to the sequencing requirements. Finally, the purified amplicons were pooled in equimolar and paired-end sequenced on an Illumina MiSeq PE250 platform (Illumina, Inc., San Diego, CA, USA) according to the standard protocols. The sequencing data obtained in this study were deposited in the NCBI Sequence Read Archive (SRA) under accession numbers SRR6201671 to SRR6201688.

### Processing of Sequencing Data

The sequences were analysed using the Quantitative Insights into Microbial Ecology (QIIME) V1.8 pipeline [15]. Then removed the average mass value of less than 20 Reads and the base number of Reads containing N over 3 Reads, the Reads length range was 220~500 nt, used the usearch to identied and removed Chimeric sequences [16]. Usearch [17] was used to cluster under the 0.97 similarity, and the OTU for species classification was obtained after clustering the sequence. Finally, all of the Clean Reads from the OTU sequence were compared, and the extracts from the reads of the OTU and the final Mapped Reads were obtained.

Richness estimates, alpha-diversity indices and beta diversity for the OTU classification were conducted using bacteria community comparisons [18]. Finally, the PCoA plots and hierarchical dendrogram based on the weighted UniFrac distance matrices [19].

### Statistical Analysis

The data of growth performance, metabolism of energy and nitrogen, serum parameters, rumen fermentation, and diversity indices were analyzed as paired t-tests (Version 9.4, SAS Institute Inc., Cary, NC, USA). Statistical significance was accepted at *P* < 0.05, and 0.05 < *P* < 0.10 was designated as a tendency. Alpha diversity indices (Chao, Simpson, Shannon) were generated with the QIIME pipeline, whereas diversity (i.e., diversity between groups of samples) was used to create principal coordinate analysis (PCoA) plots using unweighted distances. The community structure analysis histograms were produced according to taxonomy using greengenes database.

## Results

### Digestion and Metabolism of Energy and Nitrogen

Table 2 lists the results of BW, energy and nitrogen digestion and metabolism. The initial BW in the HF group did not differ from that of NF group (*P* > 0.05), but the final BW was tended to higher than that of NF group (0.05 < *P* < 0.1). The milk replacer intake in the HF group was higher than that of NF group (*P* < 0.05), however, the starter intake was tend to lower than that of NF group (0.05 < *P* < 0.1), the feed conversion rate had no differ from that of NF group (*P* > 0.05). The GE of feed intake, faecal energy (FE) and urine energy (UE) in the HF group did not differ from that of the other group (*P* > 0.05). The DE, ME, and DE/ME were higher in the NF group (0.05 < *P* < 0.1). Lambs fed HF diets had no differ apparent digestibility of GE and ME/DE compared to the NF group (*P* > 0.05). The intake N, faecal N, urine N, retained N, absorbed N, and biological values did not differ between the two groups (*P* > 0.05); however, the utilisation of N was higher than the NF group (0.05 < *P* < 0.1).

**Table 2.**
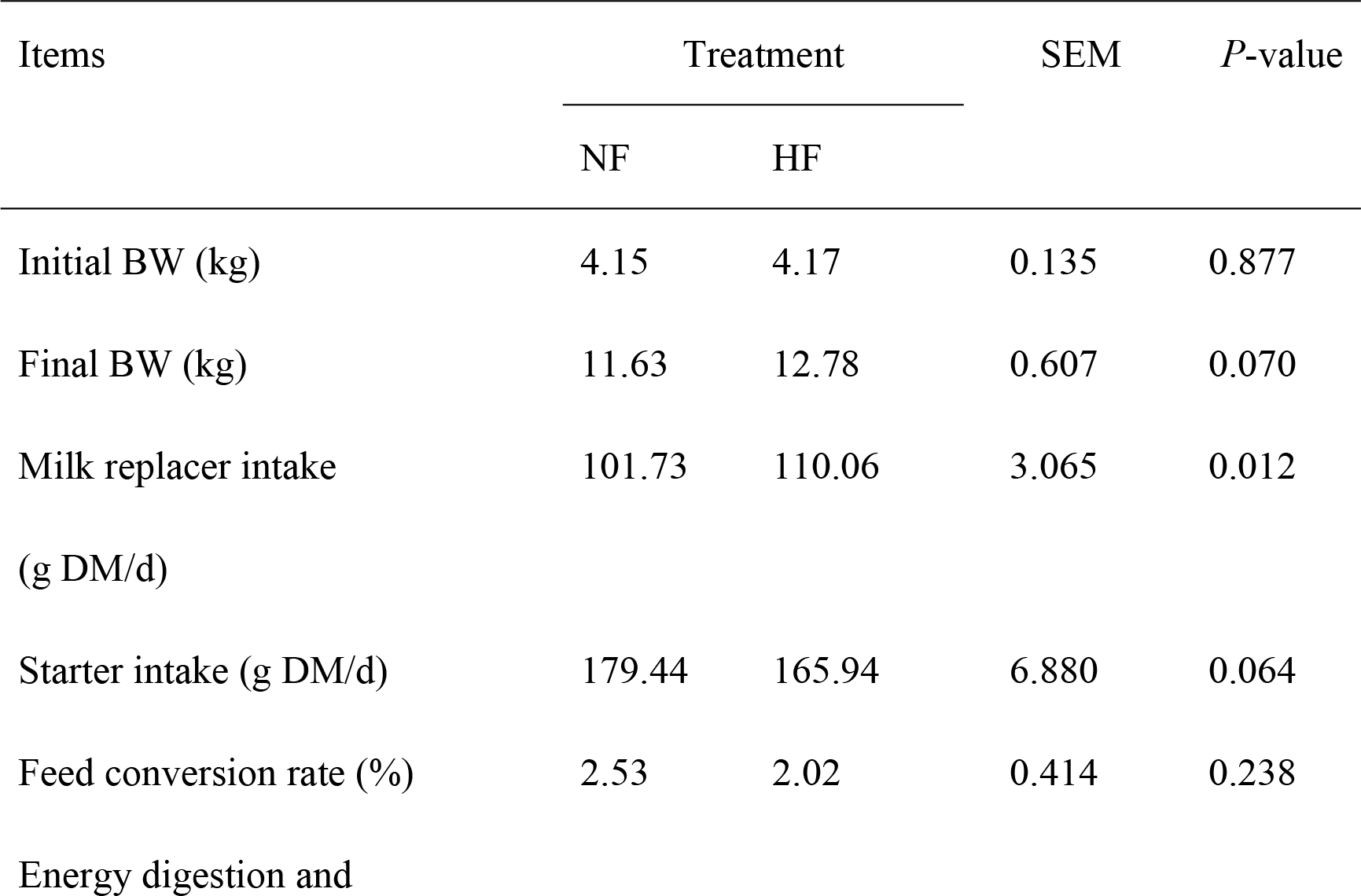
Effects of dietary fat on energy and nitrogen digestion and metabolism in twin lambs.

| Items | Treatment |  | SEM | P-value |
| --- | --- | --- | --- | --- |
|  | NF | HF |  |  |
| Initial BW (kg) | 4.15 | 4.17 | 0.135 | 0.877 |
| Final BW (kg) | 11.63 | 12.78 | 0.607 | 0.070 |
| Milk replacer intake<br>(g DM/d) | 101.73 | 110.06 | 3.065 | 0.012 |
| Starter intake (g DM/d) | 179.44 | 165.94 | 6.880 | 0.064 |
| Feed conversion rate (%) | 2.53 | 2.02 | 0.414 | 0.238 |
| Energy digestion and |  |  |  |  |

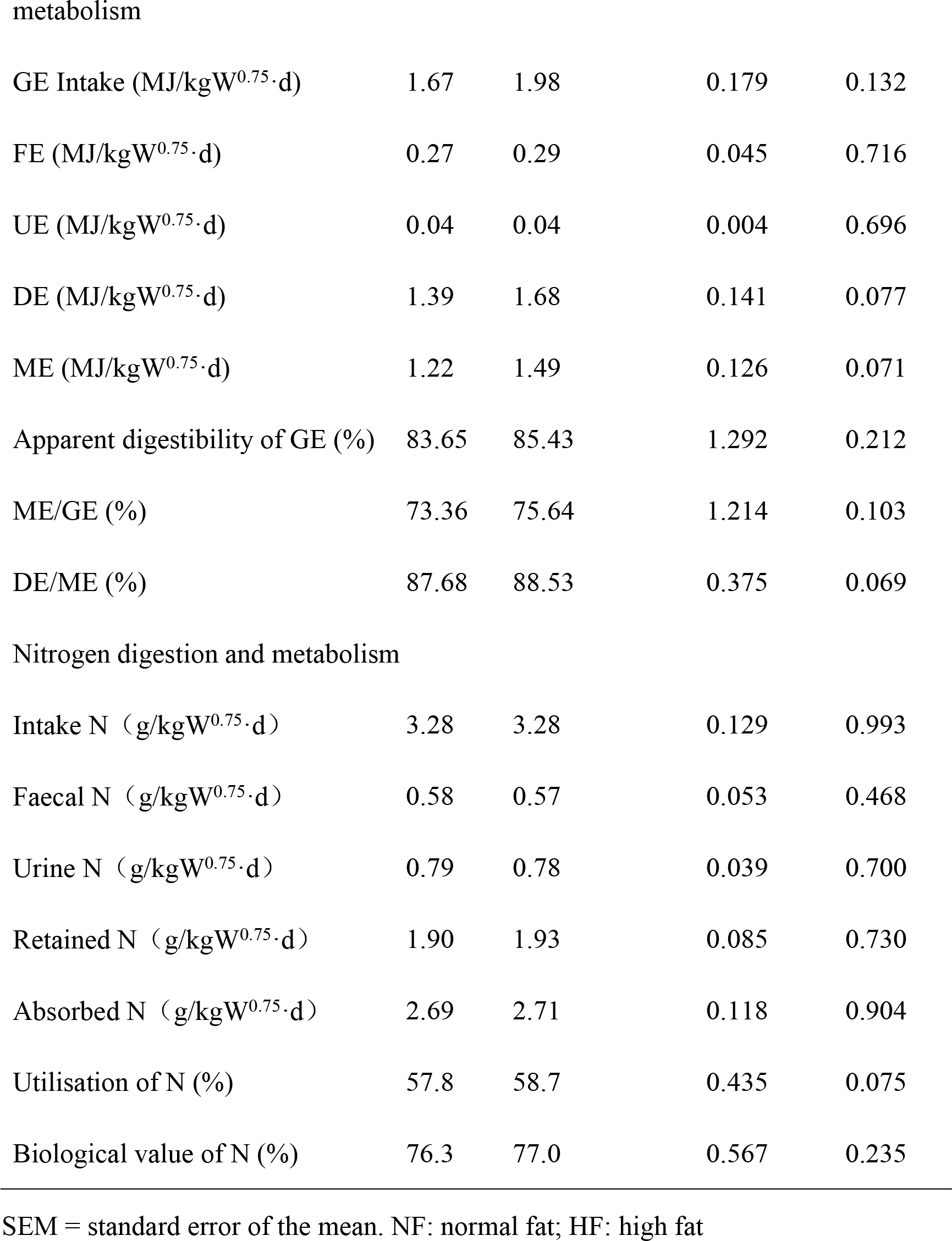

metabolism
|  |  |  |  |  |
| --- | --- | --- | --- | --- |
| GE Intake (MJ/kgW<sup>0.75</sup>·d) | 1.67 | 1.98 | 0.179 | 0.132 |
| FE (MJ/kgW<sup>0.75</sup>·d) | 0.27 | 0.29 | 0.045 | 0.716 |
| UE (MJ/kgW<sup>0.75</sup>·d) | 0.04 | 0.04 | 0.004 | 0.696 |
| DE (MJ/kgW<sup>0.75</sup>·d) | 1.39 | 1.68 | 0.141 | 0.077 |
| ME (MJ/kgW<sup>0.75</sup>·d) | 1.22 | 1.49 | 0.126 | 0.071 |
| Apparent digestibility of GE (%) | 83.65 | 85.43 | 1.292 | 0.212 |
| ME/GE (%) | 73.36 | 75.64 | 1.214 | 0.103 |
| DE/ME (%) | 87.68 | 88.53 | 0.375 | 0.069 |
| Intake N (g/kgW<sup>0.75</sup>·d) | 3.28 | 3.28 | 0.129 | 0.993 |
| Faecal N (g/kgW<sup>0.75</sup>·d) | 0.58 | 0.57 | 0.053 | 0.468 |
| Urine N (g/kgW<sup>0.75</sup>·d) | 0.79 | 0.78 | 0.039 | 0.700 |
| Retained N (g/kgW<sup>0.75</sup>·d) | 1.90 | 1.93 | 0.085 | 0.730 |
| Absorbed N (g/kgW<sup>0.75</sup>·d) | 2.69 | 2.71 | 0.118 | 0.904 |
| Utilisation of N (%) | 57.8 | 58.7 | 0.435 | 0.075 |
| Biological value of N (%) | 76.3 | 77.0 | 0.567 | 0.235 |

189 SEM = standard error of the mean. NF: normal fat; HF: high fat

### Serum Enzyme and Hormone Index

Table 3 presents the effects of dietary treatments on the serum enzyme and hormone index. No marked differences were detected between the treatments for the serum growth hormone (GH), insulin-like growth factor-1 (IGF-1), insulin (INS), leptin (LEP), adiponectin (ADP), hormone sensitive lipase (HSL), fatty acid synthase (FAS), lipoprotein lipase (LPL), and acetyl coenzyme A carboxylase (ACC) concentrations (*P* > 0.05).

**Table 3.** Effects of dietary fat on the serum enzyme and hormone index of *Hu* lambs.

| Items | Groups |  | SEM | *P*-value |
| --- | --- | --- | --- | --- |
|  | NF | HF |  |  |
| GH (ng/ml) | 1.96 | 1.72 | 0.269 | 0.179 |
| IGF-1 (ng/ml) | 23.86 | 25.95 | 3.840 | 0.357 |
| INS (uIU/ml) | 13.03 | 13.41 | 2.633 | 0.793 |
| LEP (ug/ml) | 2.97 | 3.01 | 0.097 | 0.673 |
| ADP (ug/ml) | 2.71 | 2.76 | 0.129 | 0.714 |
| HSL (U/L) | 61.12 | 63.62 | 8.301 | 0.589 |
| FAS (U/L) | 598.81 | 571.85 | 54.442 | 0.395 |
| LPL (U/L) | 71.80 | 67.01 | 4.652 | 0.132 |
| ACC (U/L) | 11.41 | 10.58 | 0.871 | 0.154 |

SEM = standard error of the mean. NF: normal fat; HF: high fat

### Rumen Fermentation Characteristics

Table 4 presents the effects of dietary fat on the characteristics of rumen fermentation. The pH values of the rumen liquid were similar (*P* > 0.05) between two groups. The concentrations of NH_3_-N and the propionate of TVFA in the HF group were higher (*P* < 0.05) than those in the NF group. The MCP production, TVFA, acetate, and acetate/ Propionate (A/P) were lower than that of NF group (*P* < 0.05). Finally, the ratio of isobutyrate, butyrate, isovalerate, and valerate did not differ in the lambs in the HF group compared with the lambs in the NF group (*P* > 0.05).

**Table 4.**
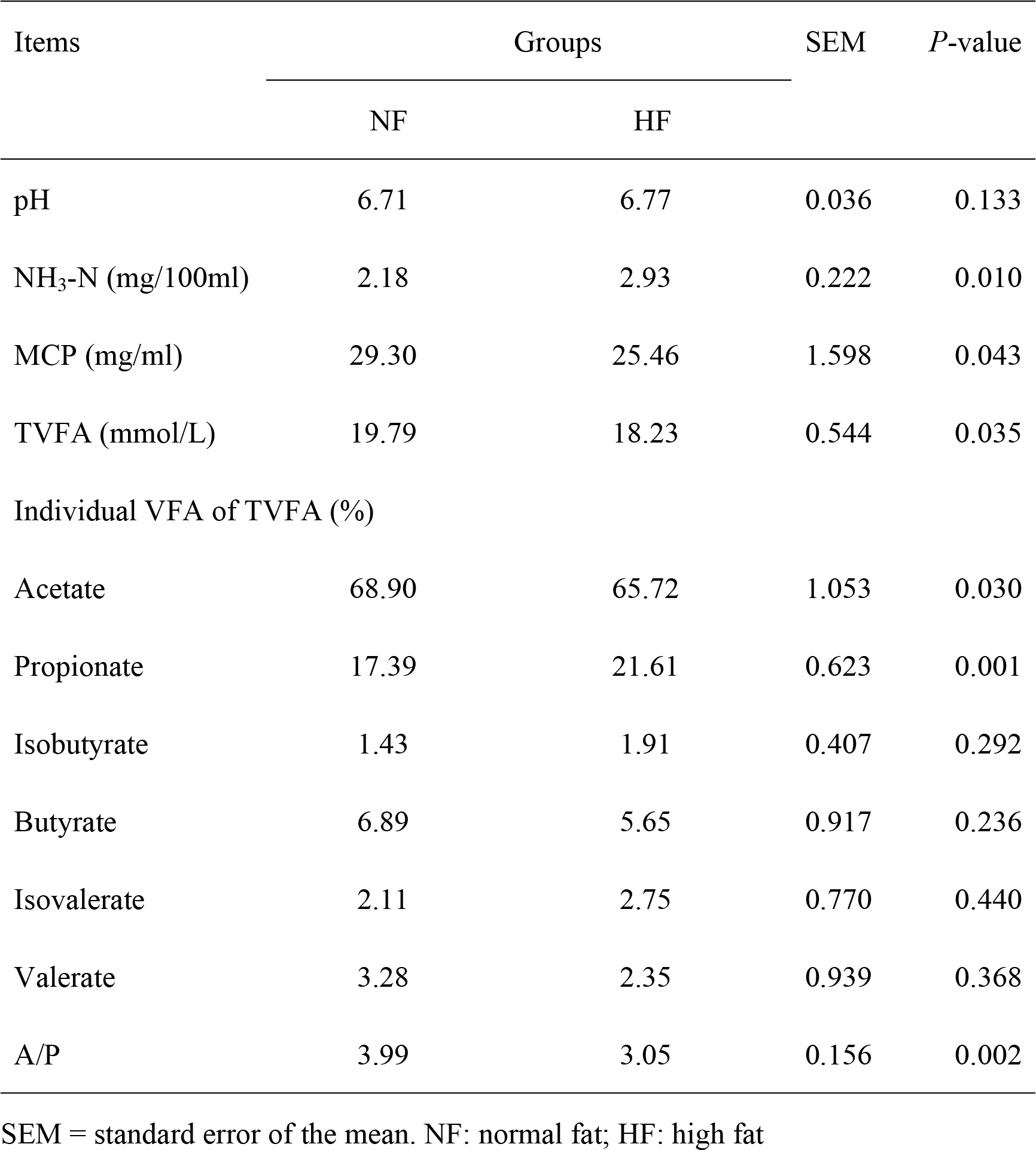
Effects of dietary fat on rumen fermentation in Hu lambs before weaning.

| Items | Groups |  | SEM | *P*-value |
| --- | --- | --- | --- | --- |
|  | NF | HF |  |  |
| pH | 6.71 | 6.77 | 0.036 | 0.133 |
| NH<sub>3</sub>-N (mg/100ml) | 2.18 | 2.93 | 0.222 | 0.010 |
| MCP (mg/ml) | 29.30 | 25.46 | 1.598 | 0.043 |
| TVFA (mmol/L) | 19.79 | 18.23 | 0.544 | 0.035 |
| Individual VFA of TVFA (%) |  |  |  |  |
| Acetate | 68.90 | 65.72 | 1.053 | 0.030 |
| Propionate | 17.39 | 21.61 | 0.623 | 0.001 |
| Isobutyrate | 1.43 | 1.91 | 0.407 | 0.292 |
| Butyrate | 6.89 | 5.65 | 0.917 | 0.236 |
| Isovalerate | 2.11 | 2.75 | 0.770 | 0.440 |
| Valerate | 3.28 | 2.35 | 0.939 | 0.368 |
| A/P | 3.99 | 3.05 | 0.156 | 0.002 |

SEM = standard error of the mean. NF: normal fat; HF: high fat

### Rumen Bacteria Composition in Different Dietary Fat Groups

S1 Table presents the number of reads obtained for each sample after the quality screening. Previous researchers have suggested that low sequencing depth leads to biased estimations of complexity curves [20]. After quality control, combining paired end reads, clustering unique and similar sequences, and filtering chimeras, 619,507 high-quality sequences from 18 samples were retained with an average of 34,417 sequences per sample. The sequencing depth was not affected by any main effect and ranged from 27,762-60,937. These sequences were assigned to 579 operational taxonomic units (OTUs) of rumen bacteria based on a 97% similarity cut-off. Furthermore, the average length of the sequence reads after primer removal was 415 bp. The Venn diagram in S1 Fig shows that the two groups shared 366 OTUs, and the NF and HF groups had 127 and 86 OTUs, respectively. There were no differences between the two groups in terms of the richness estimates and diversity indices (*P* > 0.05) (Fig. 1, Table 5).

**Table 5.**
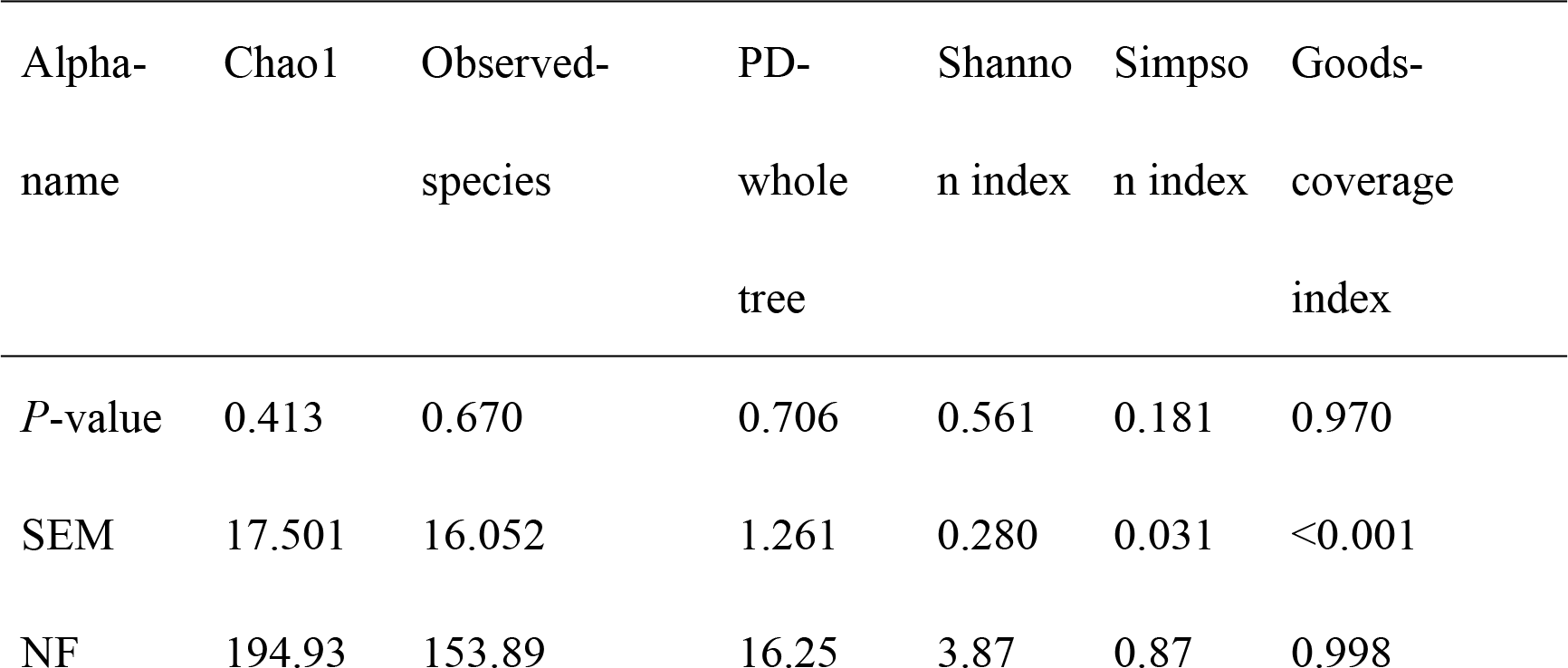
Rumen bacteria alpha diversity index in different dietary fat groups.

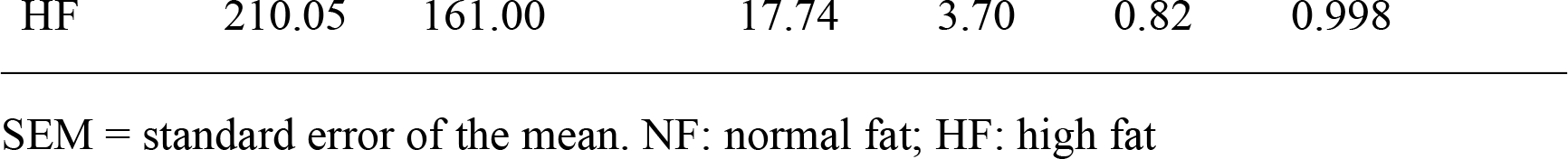

**Fig. 1:**
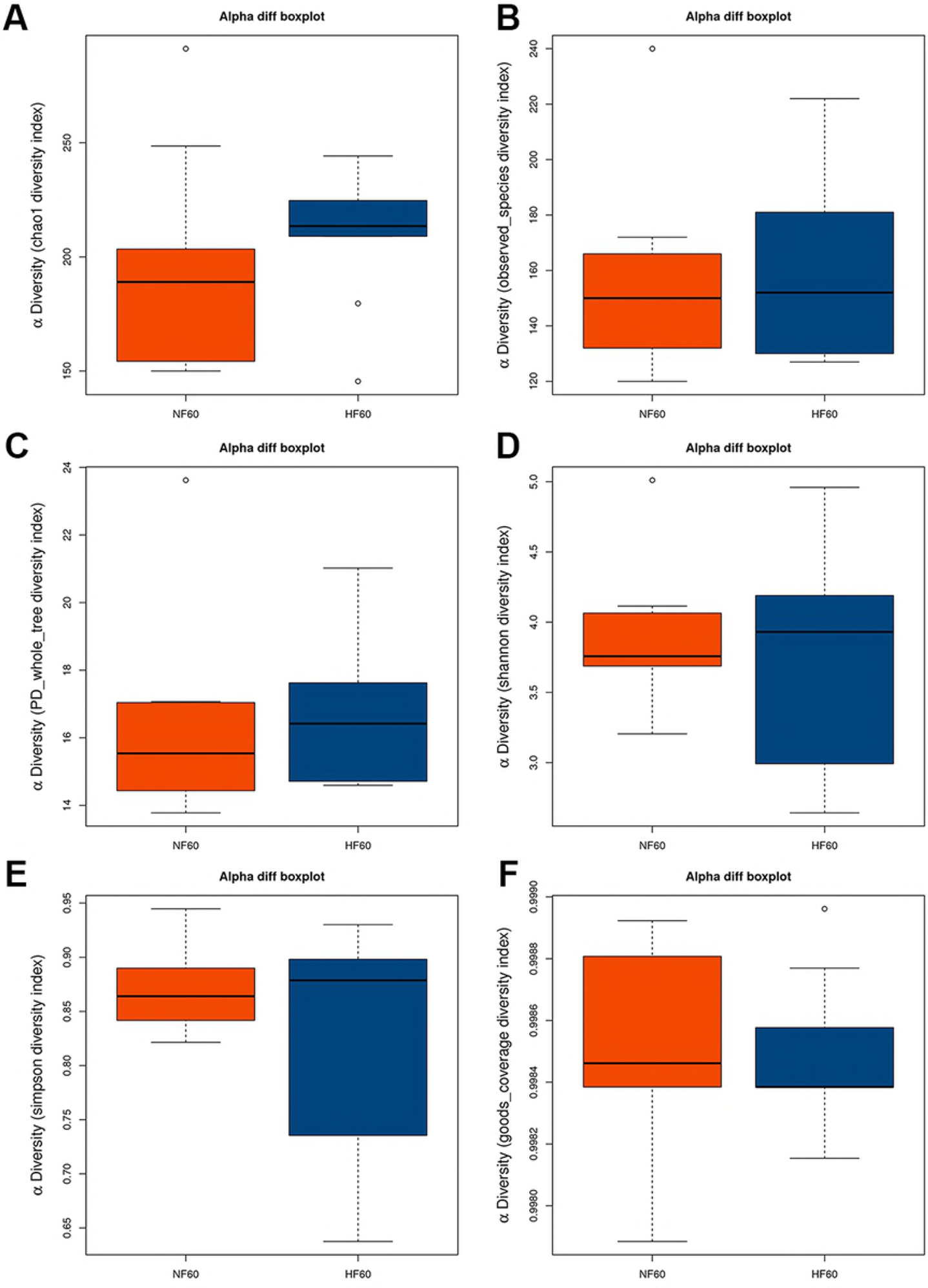
Alpha diversity index for different dietary fat groups. (A) Chao1 diversity index; (B) Observed-species index; (C) PD-whole tree index; (D) Shannon index; (E) Simpson index; and (F) goods-coverage index.

Next, we analysed the beta diversity. Fig. 2 presents the principal coordinate analysis (PCoA) with two colours that represent different dietary fat groups. The PCoA plots and hierarchical dendrogram based on the weighted UniFrac distance matrices shows that samples retrieved from different dietary fat. The PCoA plots of bacterial 16S rRNA showed no obvious clusters between the two groups that used PCoA1(45.02%) and PCoA2(18.61%). The anosim also confirmed that there were no differences (R = 0.068, *P* = 0.113, S2 Fig) between the lambs in the HF group and those in the HF group. These results confirmed that the differences between these groups were greater than those within the groups, but there was no difference between the two groups in terms of the microbial communities.

**Fig. 2:**
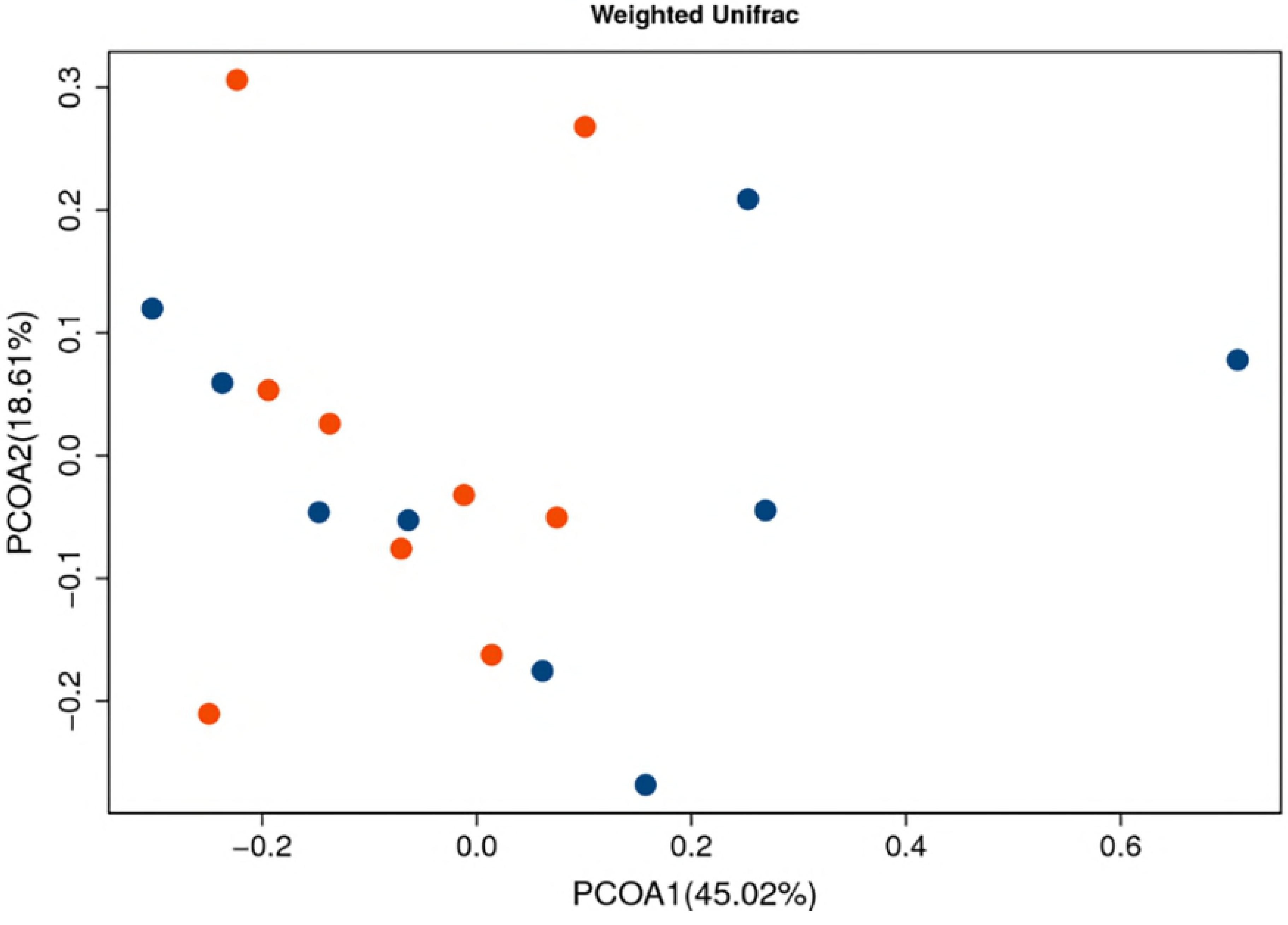
PCoA of the dissimilarity between the microbial samples. The PCoA reveals the effect of different dietary fat in twin lambs. The figures were constructed using weighted UniFrac distances.

### Taxonomic Composition of the Ruminal Bacterial Community

Fig. 3 presents the microbial compositions at the phylum and genus levels based on the taxonomic guidelines of the Silva project [21].

**Fig. 3:**
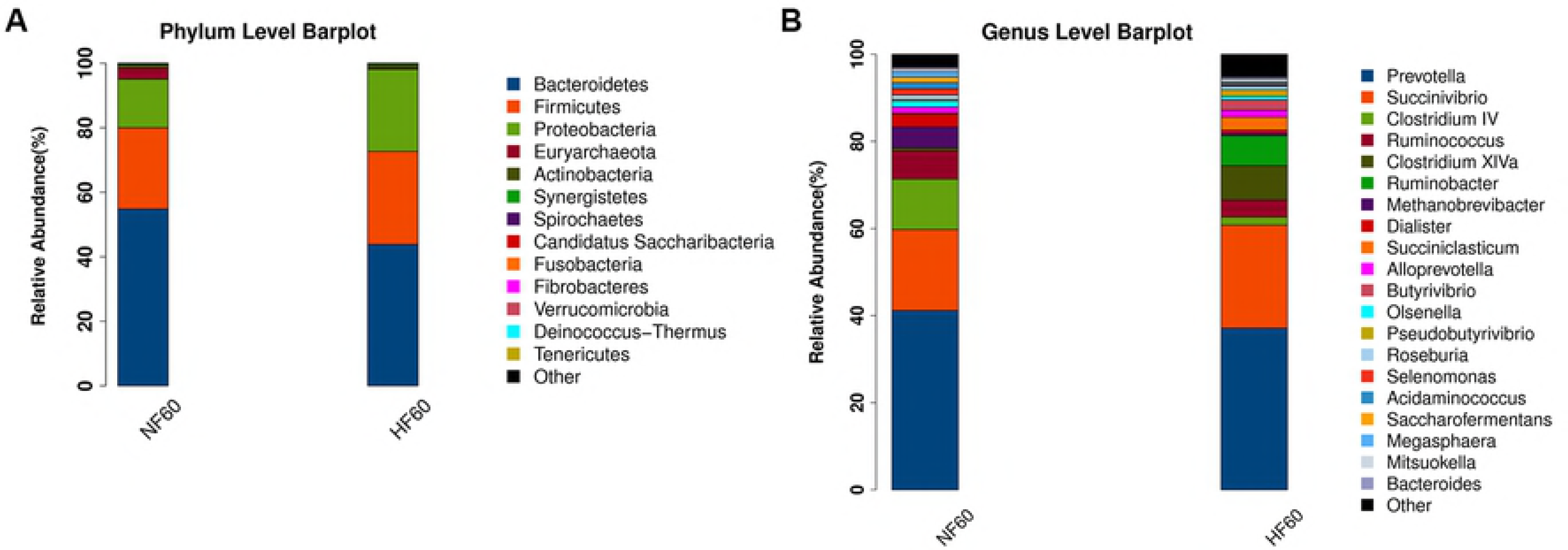
Taxonomic classification of two groups at the phylum and genus levels.(A) The relative abundance of phyla in the ruminal bacterial community and (B) the relative abundance of various genera in the ruminal bacterial community.

The first 10 phyla were detected in the rumen content samples, including *Bacteroidetes*, *Firmicutes*, *Proteobacteria*, *Euryarchaeota*, *Actinobacteria*, *Synergistetes*, *Spirochaetes*, *Candidatus Saccharibacteria*, *Fibrobacteres*, and *Verrucomicrobia*. *Bacteroidetes*, *Firmicutes* and *Proteobacteria* were identified as the dominant phyla (Table 6) in HF and NF groups. The *Bacteroidetes* phylum was the most abundant (NF 55.146% and HF 52.705%), although there were no significant differences between the two groups (*P* > 0.05). However, the *Proteobacteria* and *Fibrobacteres* phyla were higher than that in NF group (*P*<0.05). Other phyla were no significant differences (*P* > 0.05).

**Table 6.**
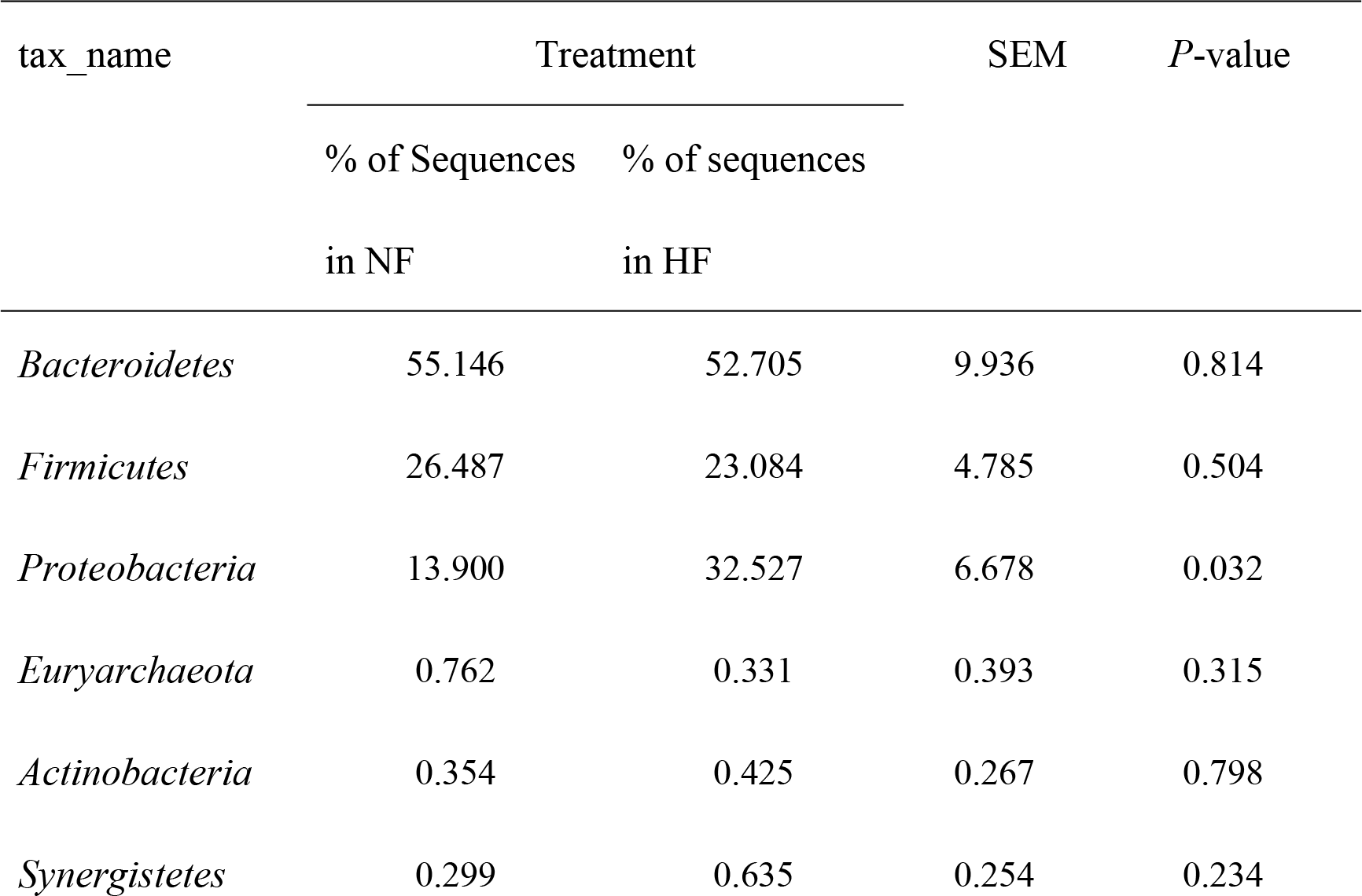
Phylum level and taxonomic composition of bacterial communities in the ruminal contents between treatment.

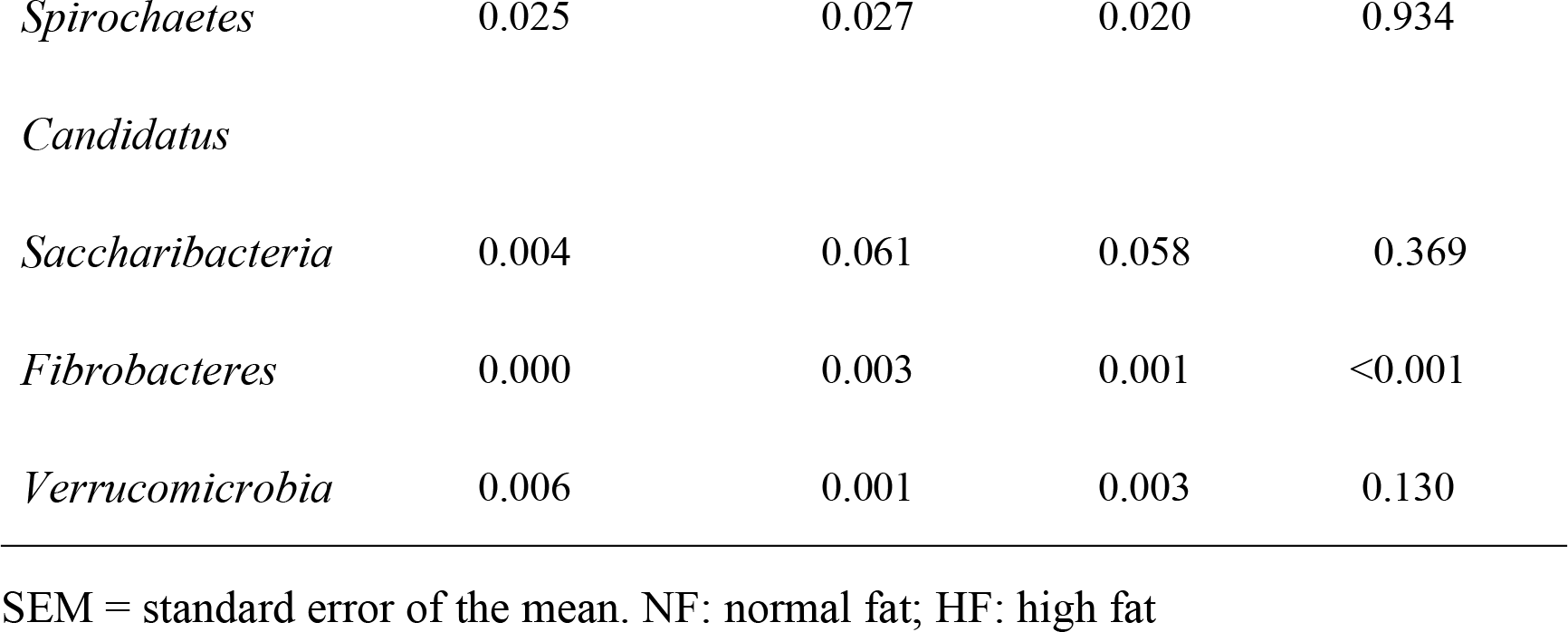

Bacterial genera containing at least 1% relative abundance are shown as a heatmap in S3 Fig. The first 20 genera identified were *Prevotella*, *Succinivibrio*, *Clostridium IV*, *Ruminococcus*, *Clostridium XlVa*, *Ruminobacter*, *Methanobrevibacter*, *Dialister*, *Roseburia*, *Succiniclasticum*, *Olsenella*, *Selenomonas*, *Butyrivibrio*, *Acidaminococcus*, *Megasphaera*, *Mitsuokella*, *Pseudobutyrivibrio*, *Alloprevotella*, *Saccharofermentans* and *Bacteroides*. The most abundant genera were *Prevotella* (40.877%), *Succinivibrio* (25.943%), *Clostridium IV* (8.579%), and *Ruminococcus* (3.694%).

At the genus level, within the *Bacteroidetes* phylum, the abundance of genus *Alloprevotella*, the genus *Succinivibrio* within the phylum *Proteobacteria*, within the *Firmicutes* phylum, the genus *Saccharofermentans* in HF group were significantly higher than that in NF group (*P* < 0.05). However, the abundance of genus *Clostridium IV*, *Dialister Roseburia*, *Butyrivibrio*, and *Acidaminococcus*, and within the *Euryarchaeota* phylum, the genus *Megasphaera* was lower than in the NF group (*P* < 0.05). There were no significant differences between the two groups in terms of the abundance of *Prevotella*, *Bacteroides*, *Ruminococcus*, *Clostridium XlVa*, *Ruminobacter*, *Methanobrevibacter*, *Succiniclasticum*, *Olsenella*, *Selenomonas*, *Mitsuokella*, and *Pseudobutyrivibrio* (Table 7; *P* > 0.05).

**Table 7.**
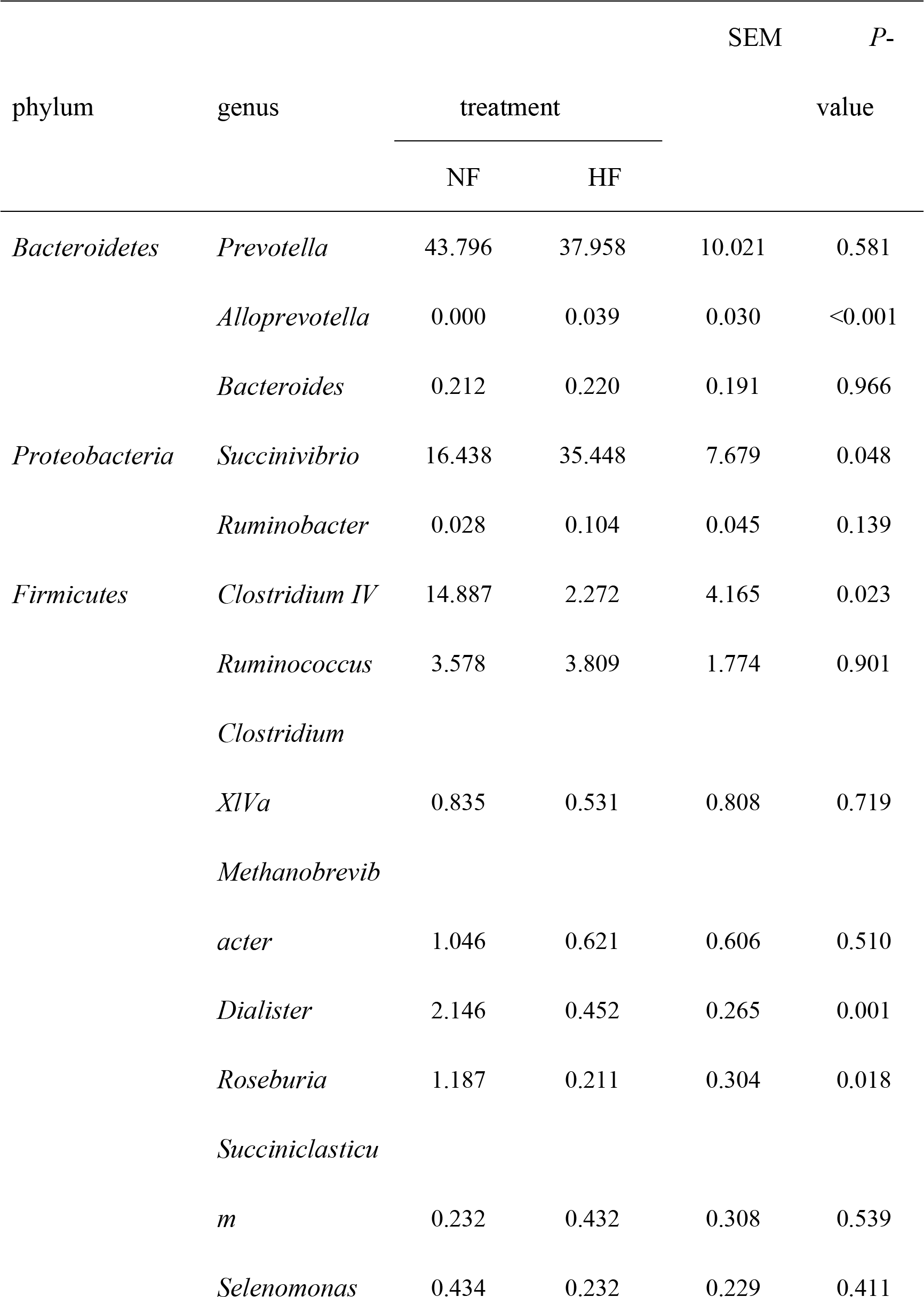
The effects of dietary fat on the relative abundance in the ruminal bacterial community.

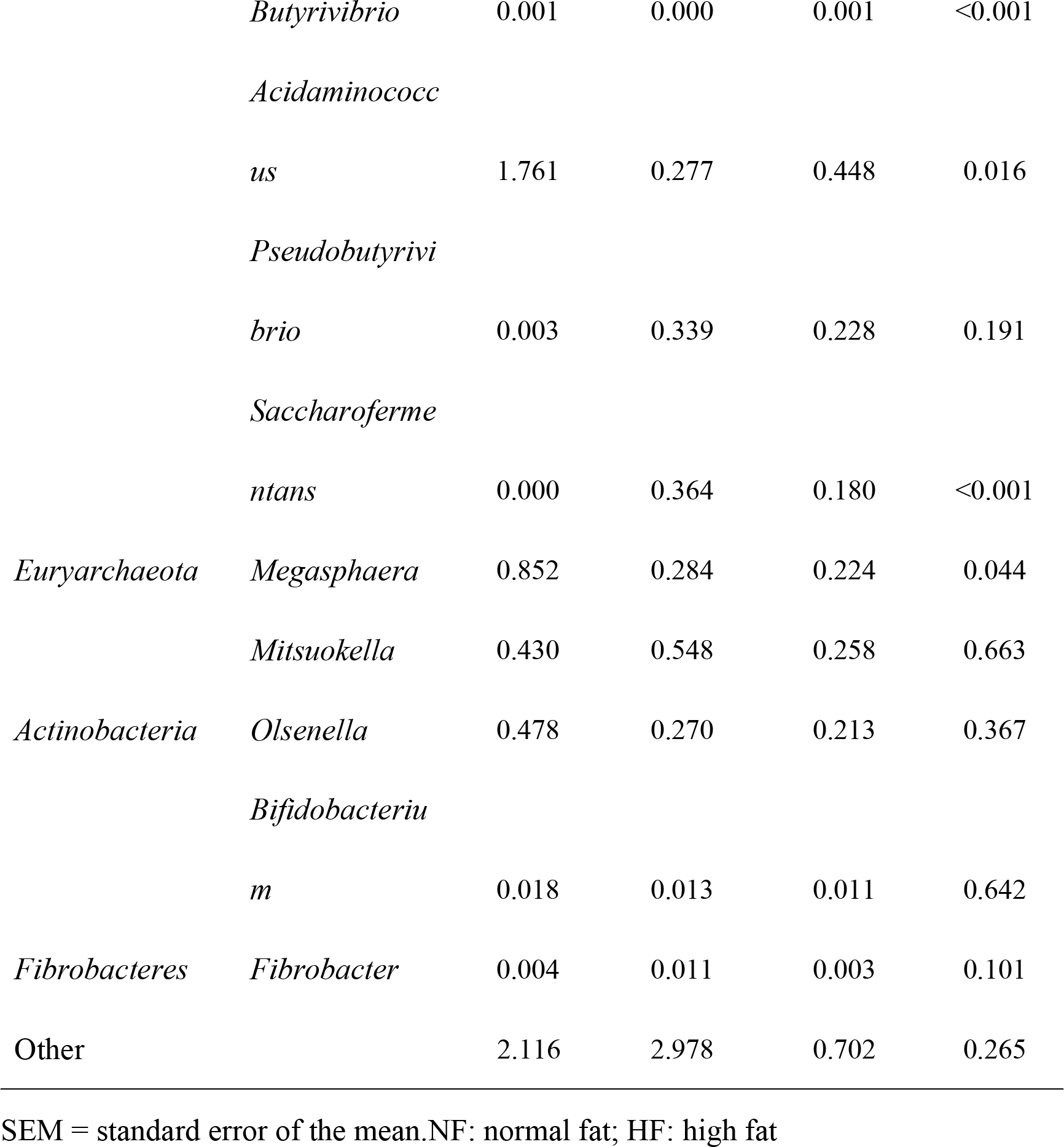

### Correlation analysis

This study evaluates the relationships between the ruminal fermentation parameter and genus abundance (the 20 most abundant shared genera; Fig. 4). The results show that the concentration of TVFA correlates positively with the abundance of *Dialister* (r= 0.500; *P*= 0.035) and *Acidaminococcus* (r= 0.527; *P*= 0.024). Meanwhile, the MCP correlates negatively with the abundance of *Ruminobacter* (r= 0.609; *P*= 0.007) and *Succiniclasticum* (r= 0.581; *P*= 0.011) and positively with the abundance of *Methanobrevibacter* (r= 0.569; *P* = 0.014), *Dialister* (r = 0.765; *P* = 0.000), *Olsenella* (r= 0.538; *P*= 0.023) and *Megasphaera* (r = 0.469; *P* = 0.049).

**Fig. 4:**
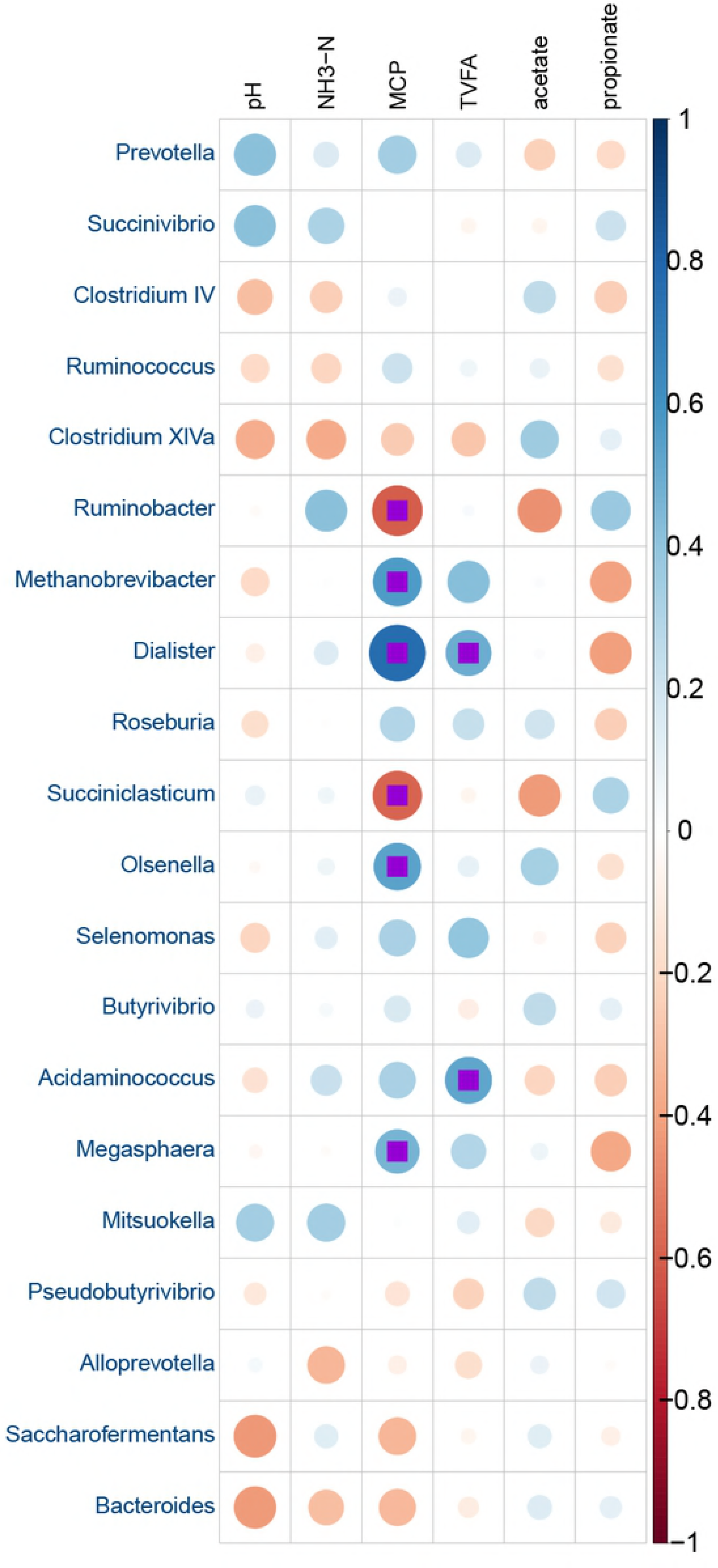
Relationships between the ruminal fermentation parameter and genus abundance. The strength of the correlation between each pair of variables is indicated by diameter and colour intensity of the circles. The square indicates a significant correlation. A colour code of dark blue indicates a positive correlation coefficient close to +1 and a colour code of dark red indicates a negative correlation coefficient close to −1.

Six taxa displayed significant differences in their abundance levels between the NF and HF groups, using LDA score log10 > 2.0 (Fig. 5A). The liner discriminant analysis effect size (LEfSe) method identified a further comparison in the NF group. These included bacteria in the genus *Methanobrevibacter* of the phylum *Euryarchaeota*; in the order *Methanobacteria* of the phylum *Euryarchaeota*; in the class *Methanobacteria* of the phylum *Euryarchaeota*; in the family *Methanobacteriaceae* of the phylum *Euryarchaeota*; and in the genuses *Dialister* and *Allisonella* of the phylum *Firmicutes* (Fig. 5B). Our results show that the relative abundance of *Methanobrevibacter*, *Dialister*, and *Allisonella* decreases significantly by improving dietary fat (Fig. 5C, 5D and 5E). S4 Fig presents the PCoA analysis of the differential species. Our findings indicate that high dietary fat may significantly affect rumen microbiota by decreasing the abundance of certain potential pathogens.

**Fig. 5:**
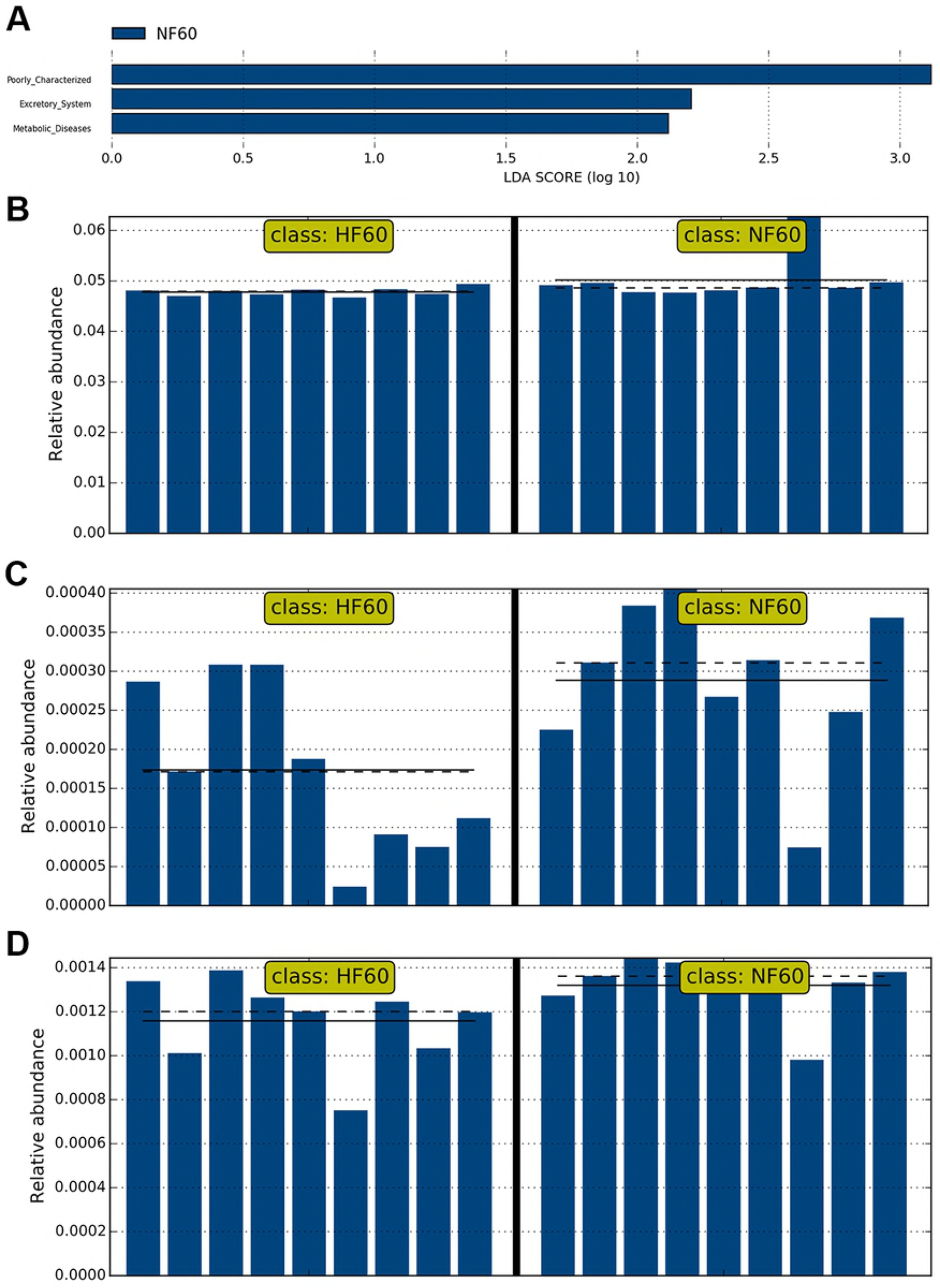
LEfSe is used to identify the most differentially abundant taxa in NF and HF samples. (A) Six taxa that meet the significant LDA threshold value of > 2 are shown in the NF and HF groups; (B) a taxonomic cladogram is obtained using a LEfSe analysis of the 16S sequences; (C) Methanobrevibacter genus; (D) Dialister genus; (E) Allisonella genus. NB: The full line indicates the group’s mean abundance value. The dotted line indicates the median of the group.

Next, we inferred further functional categories from the 16S data, and then analysed the gene sequence data of the NF group in order to estimate the functional potential of high fat diet. After adjusting for the copy number variations of the 16S rRNA genes, three main metabolic pathways were predicted from the OTU table generated using the QIIME closed-reference protocol. The results of this process showed that the NF diet might affect a broad range of biological functions, particularly the poorly characterised, excretory system, as well as metabolic diseases, compared to the HF group, which shows none of these potential functional (Fig. 6). More specifically, dietary NF might affect methane metabolism, glycolysis gluconeogenesis, aminoacyl tRNA biosynthesis, translation proteins, phenylalanine tyrosine and tryptophan, arginine and proline metabolism, valine leucine and isoleucine biosynthesis, lysine biosynthesis, streptomycin biosynthesis, RNA polymerase, bacterial toxins, base excision repair, tyrosine metabolism, phenylalanine metabolism, proximal tubule bicarbonate reclamation, protein processing in the endoplasmic reticulum, non-homologous end joining, type II diabetes mellitus, secondary bile acid biosynthesis, proteasome, retinol metabolism, primary bile acid biosynthesis, metabolism of xenobiotics by cytochrome P450, drug metabolism cytochrome P450, inorganic ion transport and metabolism and flavonoid biosynthesis. However, geraniol degradation, the vibrio cholerae pathogenic cycle, amino acid metabolism, cell motility and secretion, phenylpropanoid biosynthesis, cyanoamino acid metabolism, nitrogen metabolism, steroid hormone biosynthesis, photosynthesis, photosynthesis proteins, secretion system and bacterial invasion of epithelial cells are significantly enhanced by dietary HF (S5 Fig).

**Fig. 6:**
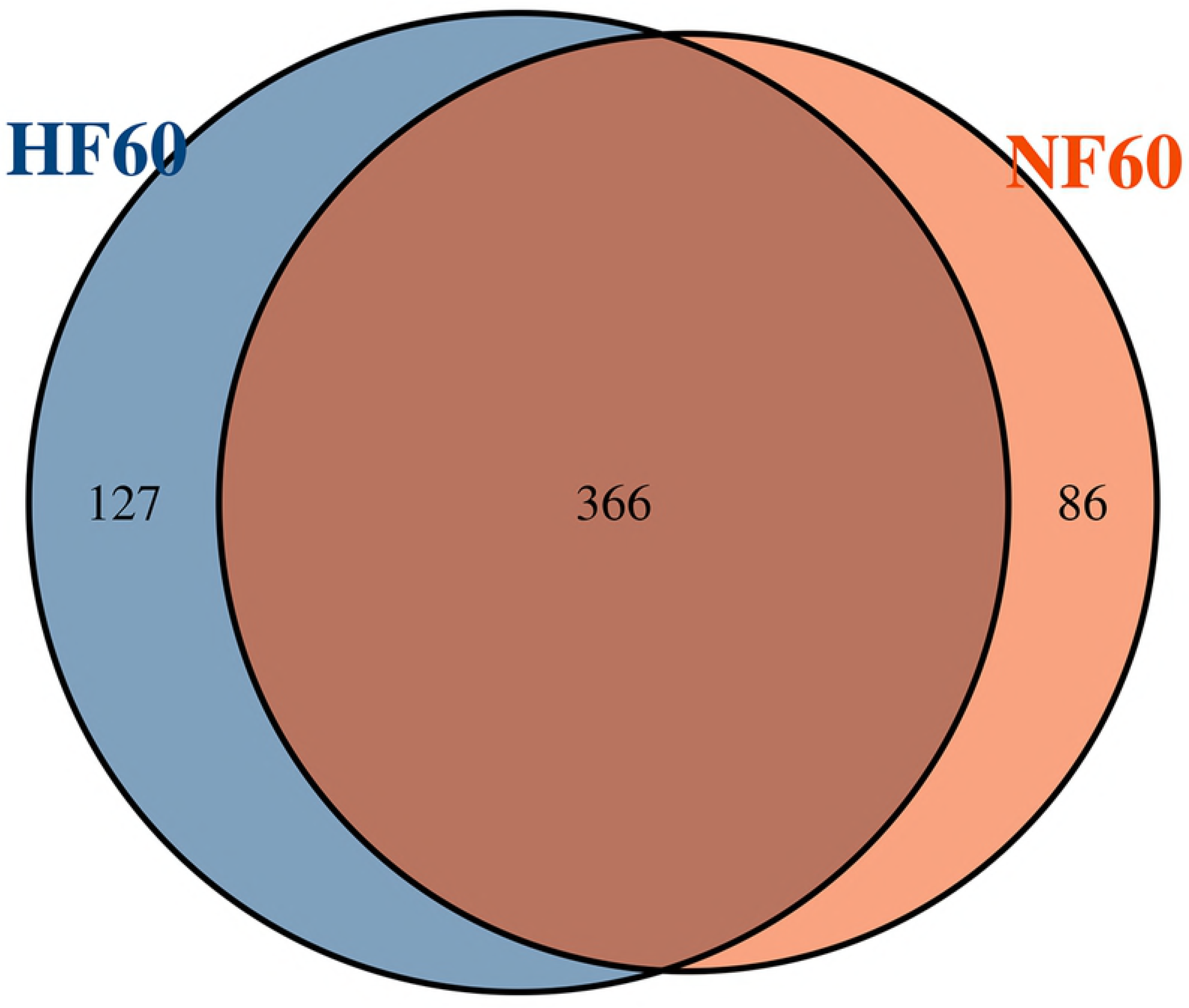
Dietary fat affects the biological pathways and functional categories of rumen microbiota.(A) Three taxa meet a significant LDA threshold value of > 2, as shown in the NF groups; (B) poorly characterised; (C) excretory system; (D) metabolic diseases. The straight line represents the mean abundance value of the group, and the dotted line represents the median of the group.

## Discussion

In the present study, the HF diet had a higher DE, ME, DE / ME, and utilisation of N. HF diet had higher fat but lower NDF in the starter, and it might slow the rumen chyme through the digestive tract, and also underdeveloped rumen has lower degradation rate and digesta passage rate of fat, which increases the digestion and absorption of other nutrients, and it finally increased higher DE, ME, and DE/ME. Therefore, dietary fat helps to improve the duodenal digestion and absorption of carbohydrates and proteins [22]. These findings are consistent with previous reports [23] that show the HF diet can improve digestion and metabolism.

FAS and ACC are key enzymes in fat synthesis, and HSL and LPL are involved in lipolysis. HGH and INS can reduce the activity of FAS, while LEP and ADP are important adipokines that are secreted by mature adipocytes [24]. These enzymes and hormones work together and ultimately determine the body fat content of animals [25]. The concentrations of FAS, ACC, HSL, INS, ADP, LEP and LPL were similar among the dietary treatments used in this study. This is not consistent with Zhang’s study on cattle [26]. Our results may indicate that the HF diet does not improve body fat content. These findings are consistent with Lundsgaard‘s recent reports [27] that show the HF diet can not effect plasma Parameters.

That may be the fat in HF diet concentration in the range of animal autoregulation. Normal pH values range from 6.4-6.8 [28]. The ruminal pH in each group in the present study appears to be within the normal range (6.71-6.77), which indicates that the internal environment of the rumen is relatively static when lambs are given the HF diet. Rumen NH_3_-N is the degradation product of feed protein [29]. In our study, the HF diet increased the NH_3_-N concentration in the rumen fluid of lambs. Brokaw suggests that adding fat to the diet prevents bacteria from adhering to the feed, thereby reducing the synthesis of MCP and causing an increase in NH_3_-N [30]. The results of the current study regarding MCP are consistent with previous findings. It is known that VFA were produced by the degradation of carbohydrates, and its concentration is affected by many factors, such as diet composition, feeding methods, and nutrition level [31]. Shaver [4] concludes that including up to 5% of tallow in the diet has minimal negative effects on rumen fermentation. Pantoja later came to the same conclusion [32]. However, the results of our experiment were similar to that of Onetti’s, as the proportion of ruminal acetate decreased and the proportion of propionate increased using the HF diet. As a result, the A/P also decreased. Possible causes of this result could be the animal species, raw materials and type of fat used in the study.

Diet composition is an important factor that influences the function and structure of microbial communities in the rumen [33,34]. The high-throughput sequencing approach is a powerful way to reveal the bacterial diversity of rumen contents. Good’s coverage estimate in the current study was 0.998, which indicates that the research recovered more than 99% of all OTUs and computed with a similarity rate of 0.97. The diversity indices and richness estimates are similar, which indicates that the rumen microbial diversity of the HF group was similar to that of the NF group [35]. In our study, the HF and NF groups had the same levels of protein and feed ingredients, and the lambs were raised under the same environmental conditions, which might explain the similarities in the rumen microbial diversity. Moreover, the PCoA, which was used to examine phylogenetic divergence, did not significantly cluster among the OTUs, thus demonstrating further that there was no difference between the microbial communities of the two groups.

Furthermore, the rumen microbial community was complex and dependent on the microbial richness, diet structure, diet composition and physiology of the host [36,37]. *Bacteroidetes*, *Firmicutes* and *Proteobacteria* were the most abundant phylum in this study. These are the main bacterial phyla, and they play vital roles in rumen fermentation [38,39]. Jami and Mizrahi [40] reported that the proportion of *Bacteroidetes* was higher than *Firmicutes*, which is consistent with our results. Previous research shows that *Proteobacteria* is dominant in newborn lambs, followed by a sharp decline where it becomes the lowest proportion and *Bacteroidetes* becomes the highest [41,42]. In our study, the proportion of *Proteobacteria* and *Fibrobacteres* phylum was still high in HF group, which is similar to the study on the gut microbiome of mice fed with high fat diet by Hildebrandt [43]. It might be the two phylum bacteria preferring a high fat environment.

As previously reported, *Prevotella* within the *Bacteroidetes* phylum is responsible for cleaving oligopeptides and plays an important role in protein metabolism [44]. It also contributes to the majority of hereditary and metabolic varieties of the microflora [45]. Furthermore, *Prevotella* is the most abundant genus in adult rumen [46,47]. This is in agreement with our result, which show that *Prevotella* is predominantly composed of this genus when a high-caloric diet is consumed [48]. The HF treatments did not affect the relative abundance of *Prevotella*; therefore, our results are similar to those of Stevenson [44]. The genus *Succiniclasticum* of *Proteobacteria* phylum, which is the second most abundant bacteria, converts succinic acid fermentation into propionic acid and improves the bioavailability of butyrate for the host [49]. In this study, the proportion of *Succiniclasticum* in HF exceeds the NF group numerically, which may indicate some degree of resource competition among the rumen bacteria [50]. This may be one of the reasons for the increased concentration of propionic acid in the rumen of the HF group.

*Clostridium* is a genus of Gram-positive bacteria within the phylum *Firmicutes* and it includes several significant human pathogens, including the causative agent of botulism, which is a leading cause of diarrhoea [51]. The alteration of the microbiota through the HF diet in this study was inconsistent with previous reports that show an HF diet increases Clostridium, which belongs to the phylum *Firmicutes* [52]. One possible explanation is that the fat of HF diet in this study is 1.8 times higher than the fat of NF diet, and less than 60% of fat was added of the former. This result also shows that the HF diet was beneficial to lamb health under the experimental conditions.

The genus *Ruminococcus* of the phylum *Firmicutes* is a major genus that decreases numerically through the HF diet. *Ruminococcus* can produce cellulase and hemicellulose, which decomposes plant fibre [53]. The results showed that the proper increase of fat had no effect on the abundance of genus *Ruminococcus* in the rumen. *Acidaminococcus* is a genus within the phylum *Firmicutes* that produces acetic acid and butyric acid [54]. This could explain the acetic acid in the rumen of the NF group, which was significantly higher than that of the HF group. The PUFA present is believed to have a toxic effect on cellulolytic bacteria [55]. In the present study, no effect was observed on some cellulolytic genera (such as *Fibrobacter* and *Ruminococcus*), the *Butyrivibrio* also belong to cellulolytic genera in HF group is lower than that in NF group. This indicates that dietary fat at the level used in the present study does not have a toxic effect on cellulolytic bacteria. Of course, it also has a great relationship with the type of animal and the stage of growth.

Next, we explored the relationship between the rumen parameters and ruminal microbiota. The results indicate that several rumen bacterial genera were affected by dietary fat, which could be linked to changes in dietary composition, differences in ruminal fermentation characteristics. A correlation analysis revealed that there was a relationship between the proportion of rumen TVFA and the rumen microbes. The increased TVFA was associated with the genera *Dialister* and *Acidaminococcus*, which belong to the phyla *Firmicutes*. This finding suggests that these two genera might be involved in nitrogen and volatile fatty acid metabolism, as well as fibrolytic enzyme secretion and starch degradation, which is consistent with Wang's findings [37]. The abundances of the genera *Ruminobacter* and *Succiniclasticum* were each found to be negatively correlated with rumen MCP. Furthermore, the correlation analysis showed that the concentrations of rumen MCP were linked to enrichments in the abundances of the genera *Methanobrevibacter*, *Dialister*, *Olsenella* and *Megasphaera*, which indicates that they strongly adhered to feed and produced large amounts of MCP. The differential species analysis in the present study revealed three important pathways. The *Methanobrevibacter* genus utilised hydrogen to reduce carbon dioxide into methane, which caused energy loss that ranged from 2-12 % of the cattle’s gross energy intake [56]. Therefore, this study infers that the *Methanobrevibacter* genus can affect the pathway of energy metabolism, and increasing fat can reduce methane production within a certain range. *Dialister* genus within the phylum *Firmicutes* was related to lipid and glucose metabolism, glucose tolerance and the inflammatory immune responses [57,58]. This was similar to our experimental predictions. The genus *Allisonella* of the phylum *Firmicutes* caused an anti-inflammatory commensal and reduced inflammation in patients with bowel disease and irritable bowel syndrome [59]. The reduction of the *Allisonella* genus in the HF group might be associated with the inflammatory response, but the specific reasons require further research.

## CONCLUSIONS

This study demonstrated that high fat diet can affect the abundance of several groups of rumen bacteria in rumen, such as significantly increasing phyla *Proteobacteria* and *Fibrobacteres*, and genera of *Succinivibrio*, *Alloprevotella*, and *Saccharofermentans*, but significantly decreasing genera of *Clostridium IV*, *Dialister*, *Roseburia*, and *Butyrivibrio*. And high fat diet improved the performance of lambs at weigh gain and energy utilization, and had effect on VFA composition but no effects on serum enzymes and serum hormone index. Considering the potential effects on rumen fermentation, future studies are required to further investigate how the high fat diet affect the rumen function of lambs during the stage.

## Competing interest

The authors declare that they have no competing interest.

## Acknowledgements

We thank the staff (Fan Zhang, Bo Wang, and Yan. Tu) of Feed Research Institute of Chinese Academy of Agricultural Sciences for their technical assistance. The China Agriculture Research System (CARS-38) and Special Fund for Agro-scientific Research in the Public Interest (201303143) supported this study.

## List of figures and tables

**Table 1. Ingredients and nutritional composition of milk replacer and starter.**

**Table 2. Effects of dietary fat on energy and nitrogen digestion and metabolism in twin lambs.**

**Table 3. Effects of dietary fat on the serum enzyme and hormone index of Hu lambs.**

**Table 4. Effects of dietary fat on rumen fermentation in Hu lambs before weaning.**

**Table 5. Rumen bacteria alpha diversity index in different dietary fat groups.**

**Fig. 3: Taxonomic classification of two groups at the phylum and genus levels. Table 6. Phylum level and taxonomic composition of bacterial communities in the ruminal contents between treatment.**

**Fig. 4: Relationships between the ruminal fermentation parameter and genus abundance.**

**Fig. 5: LEfSe is used to identify the most differentially abundant taxa in NF and HF samples.**

**Fig. 6: Dietary fat affects the biological pathways and functional categories of rumen microbiota.**

## Additional Files

**S1 Table: Downstream analysis. doc**

**S1 Fig:**
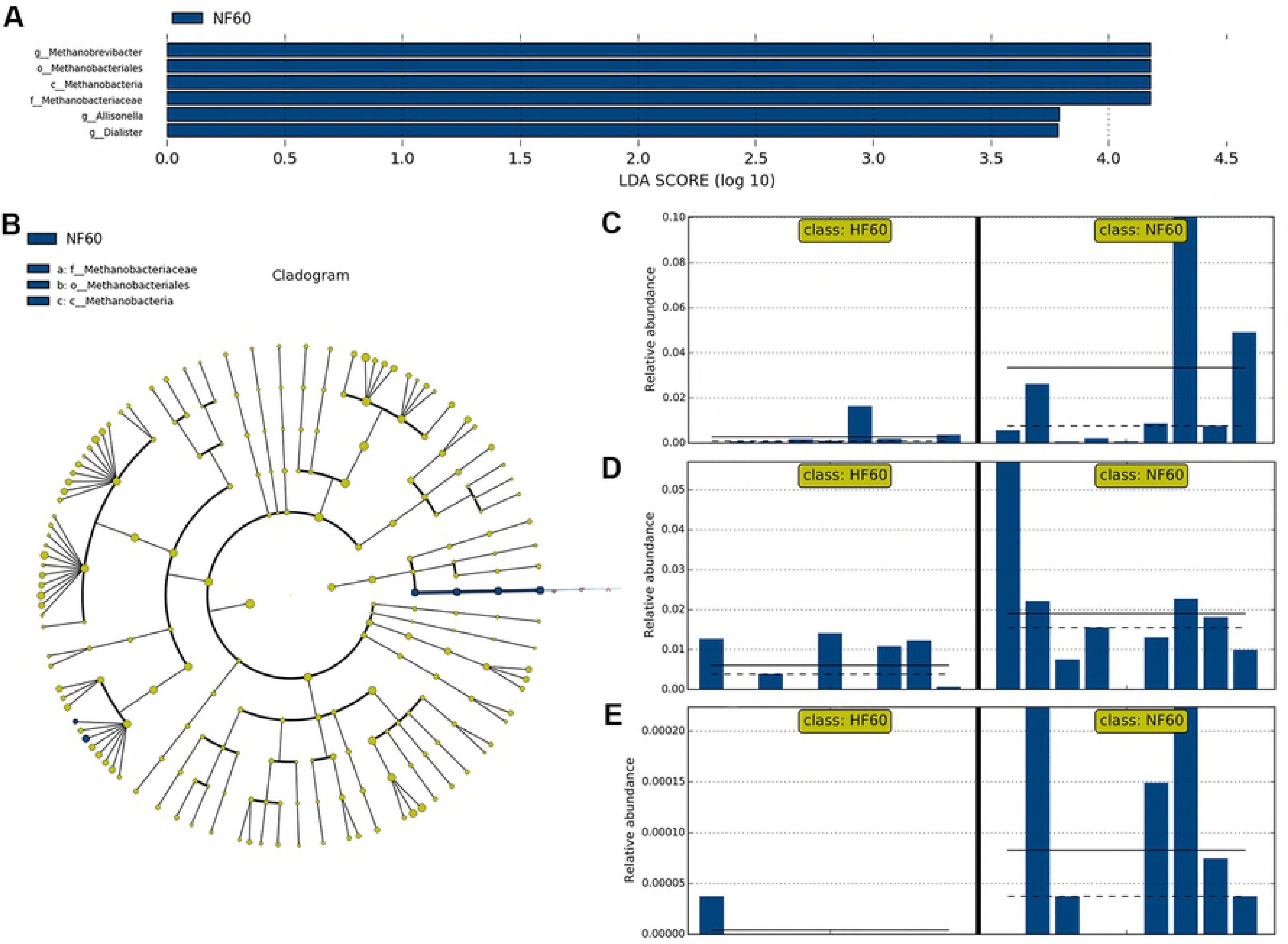
Venn diagram summarizing number of OTU between NF group and HF group. tiff.

**S2 Fig:**
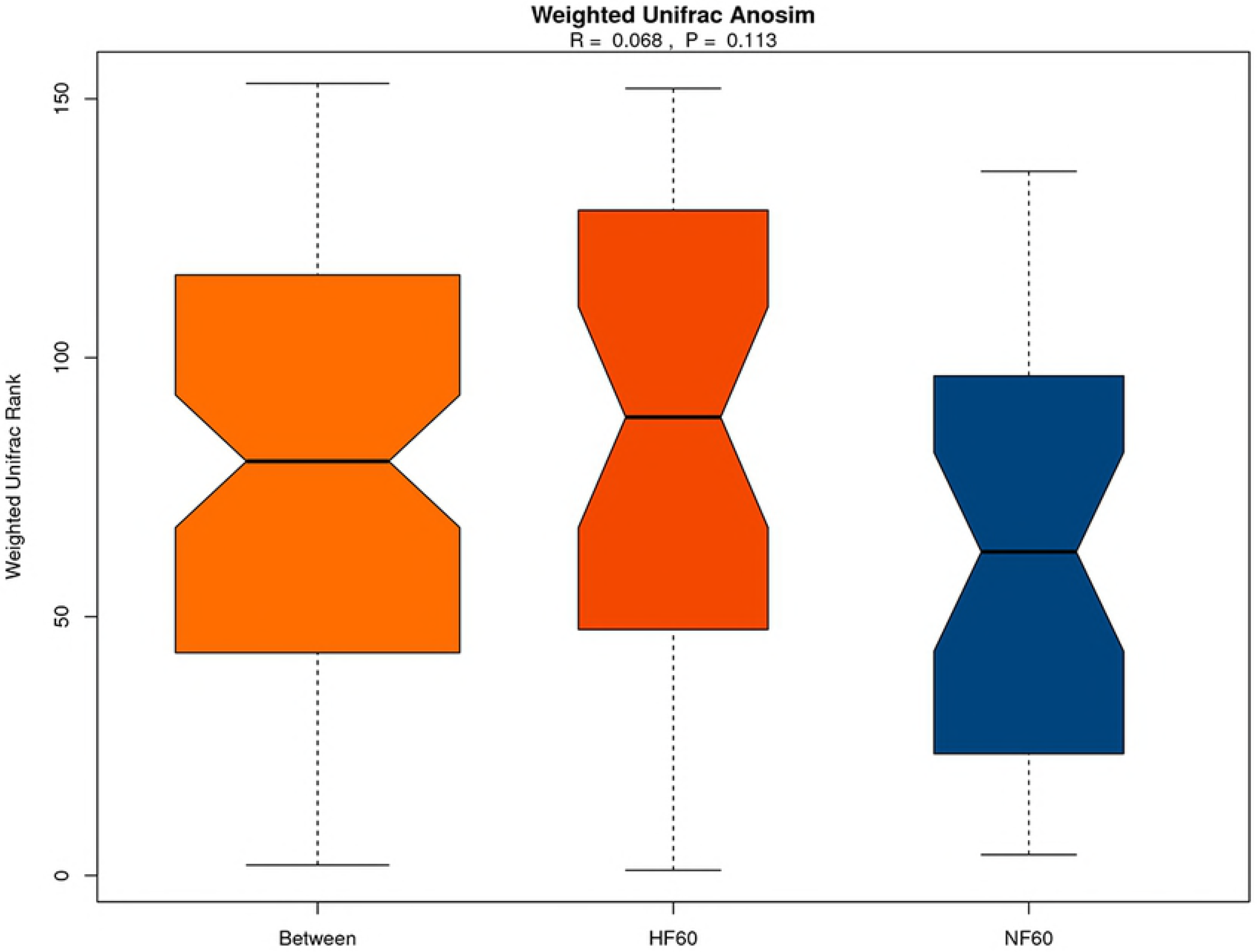
Anosim similarity analysis. Figures are constructed using weighted UniFrac distances. tiff.

**S3 Fig:**
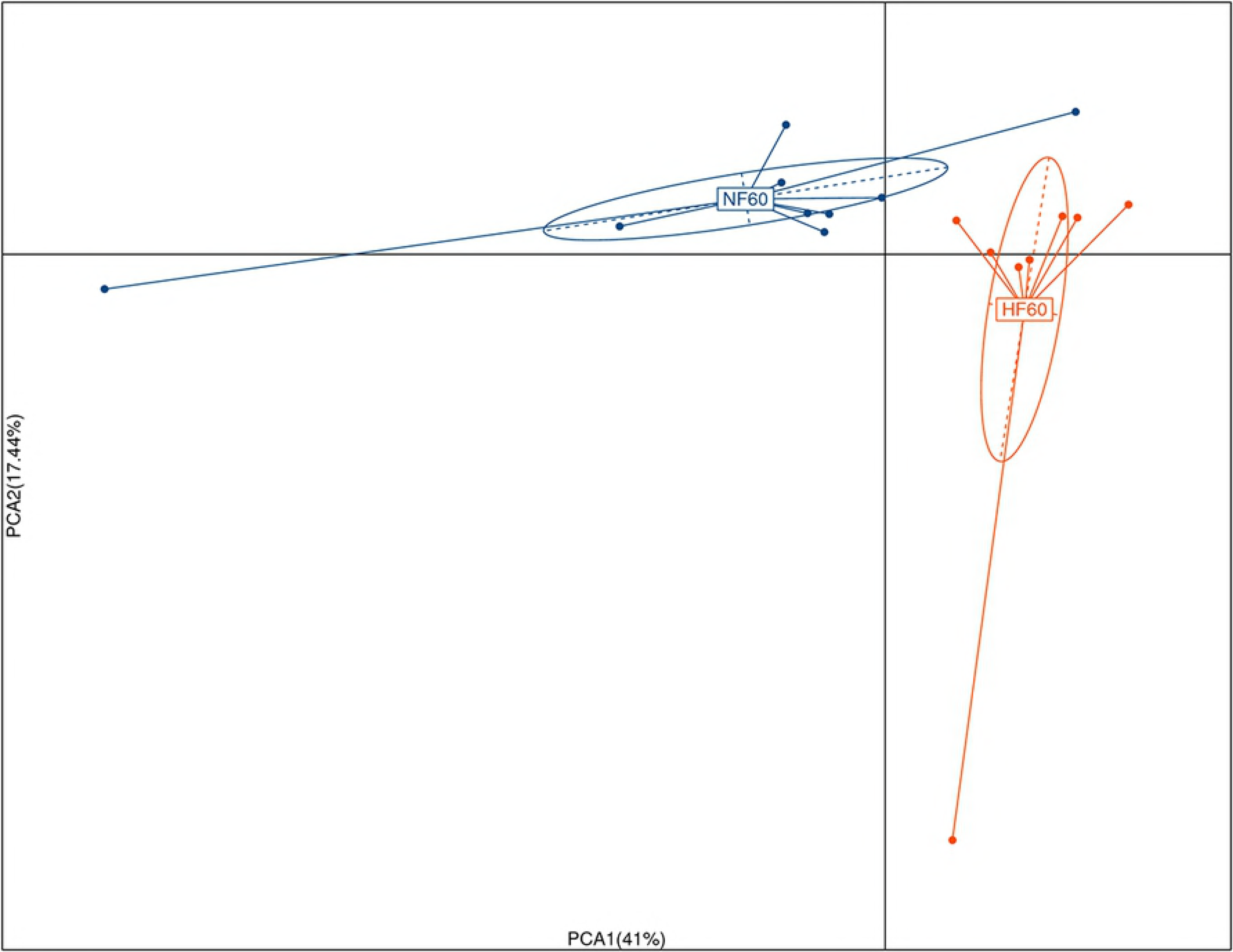
Heatmap showing the bacterial community composition based on analysis of the 30 most abundant genera. tiff.

**S4 Fig:**
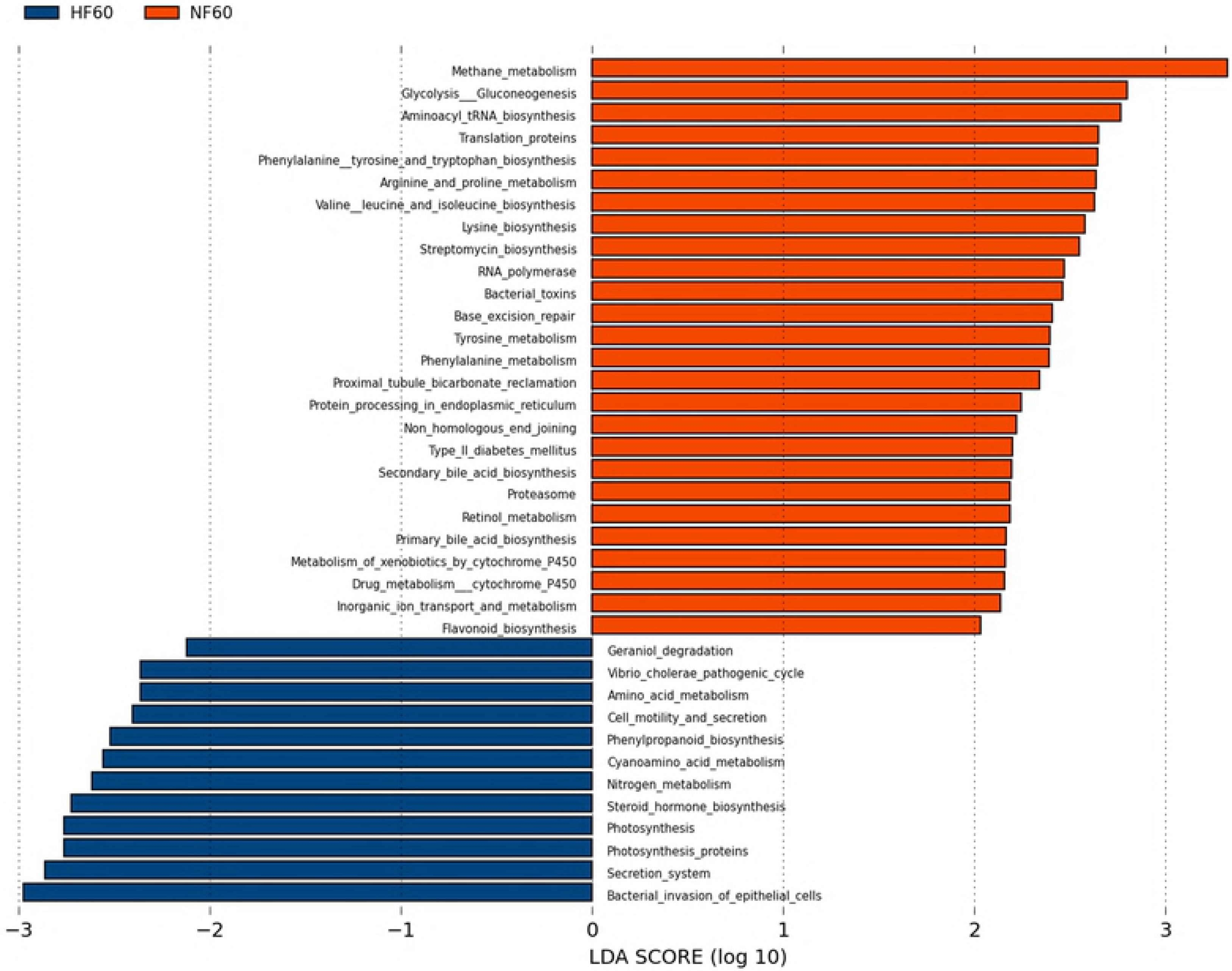
PCoA of the dissimilarity between microbial samples. Figures are constructed using weighted UniFrac distances. tiff.

**S5 Fig:**
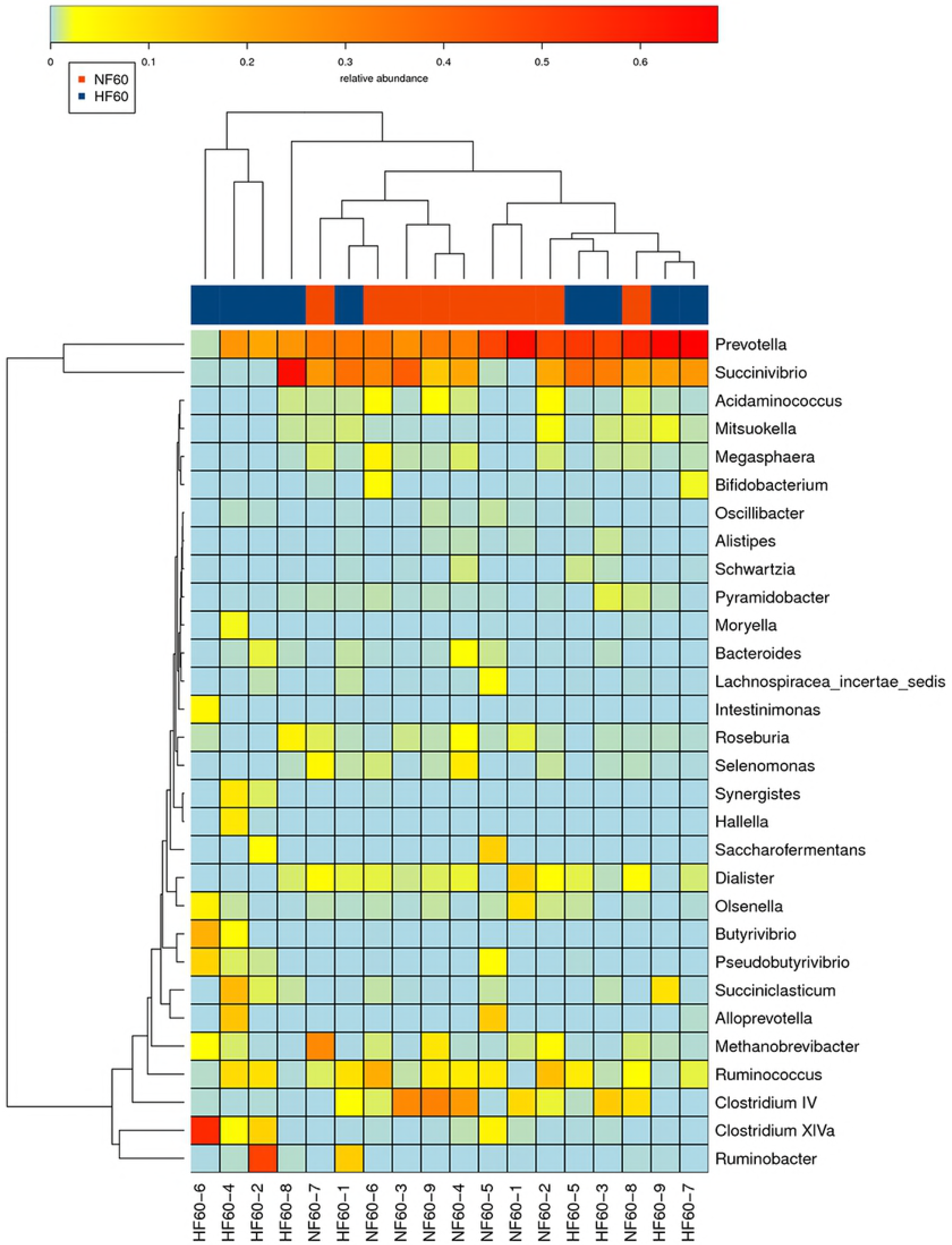
Prediction of the differential species function metagenomes of NF and HF ruminal microbial communities. tiff.

